# Sympathetic signaling directs macrophage efferocytosis in thermogenic adipose tissue

**DOI:** 10.1101/2025.10.28.685032

**Authors:** Simon Meyer, Jennifer Witt, Imke Liebold, Tamara López-López, Laura Ehlen, Stephanie Leyk, Johanna Hiefner, Daniel Haas, Christian Schlein, Pablo J. Sáez, Natalie Krahmer, Jorg Heeren, Lorenz Adlung, Lidia Bosurgi, Anna Worthmann

## Abstract

Brown adipose tissue (BAT) undergoes significant remodeling upon thermogenic activation. During this process, brown adipocytes and immune cells, such as macrophages, contribute to thermogenesis and energy expenditure. Among the various functions exerted by macrophages, the clearance of dying cells, known as *efferocytosis*, is a key regulator of tissue remodeling across multiple organs in both physiological and pathological contexts. However, whether macrophages contribute to BAT remodeling and thermogenic adaptation through efferocytosis, and what drives efferocytosis in BAT, remain unknown.

Here, we identify norepinephrine (NE), which is highly released in BAT upon cold challenge, as a tissue-specific trigger of macrophage efferocytosis. Transcriptomic and lipidomic analyses of BAT after cold exposure revealed a population of lipid-handling macrophages enriched in efferocytosis-related transcripts. Consistently, cold exposure enhanced the efferocytic capacity of BAT macrophages. These effects were recapitulated by stimulation of macrophages with NE and were dependent on β2-adrenergic signaling and the efferocytic receptors AXL and MERTK. Mice lacking *Axl* and *Mertk* in macrophages exhibited impaired lipolysis, reduced thermogenic gene expression, and increased adipose tissue inflammation.

Together, our findings identify a so far neglected role for NE in adipose tissue, linking sympathetic activation to macrophage efferocytosis and thereby promoting tissue remodeling and metabolic adaptation. Uncovering the role of NE in one of the core functions of macrophages, efferocytosis, not only expands our understanding of the multifaceted effects of NE on the immune system but also highlights therapeutic potential for targeting impaired efferocytosis in metabolic disorders.

**One sentence summary:** Norepinephrine is a novel trigger of macrophage efferocytosis in brown adipose tissue, linking sympathetic signaling to metabolic adaptation and macrophage tissue remodeling responses through β2-adrenergic and *Axl/Mertk* pathways.

## INTRODUCTION

It is estimated that every day in the human body up to 300 billion cells undergo apoptosis and are taken up by macrophages in a process called efferocytosis (*1*). Even though considered immunologically silent under homeostatic conditions, this process critically affects macrophage function and hence regulate tissue remodelling and inflammatory responses. Dysregulation of this process is implicated in the pathogenesis of various disorders including atherosclerosis and autoimmunity (*2–12*). The mechanisms controlling macrophage reprogramming through efferocytosis remain under investigation (*13*), but a deep understanding of the mediators controlling efferocytosis efficiency is crucial to harness this process for therapeutic purposes. To date, mediators that activate nuclear receptors such as LXR and PPARs, have been identified as key drivers of efferocytic capacity (*14*, *15*). Efferocytosis also varies across different tissues due to both the nature of the phagocytic cells involved and tissue-specific microenvironments (*16*). Tissue-specific signals modulating efferocytosis efficiency are thereby beginning to be elucidated. For instance, adiponectin, released by adipocytes, induces the expression of macrophages’ efferocytic receptors such as MERTK and increases macrophage efficiency to recognize and engulf apoptotic cells via engagement of the nuclear receptor retinoid X receptor (*17*).

Adipose tissue, in particular, undergoes dynamic remodeling influenced by metabolic state and may therefore be especially dependent on efferocytosis efficiency. Depending on environmental signals, energy can be released and stored especially in white adipose tissues (WAT) or used for thermogenesis in brown adipose tissues (BAT) overall strongly affecting tissue mass and composition (*18*). In the context of obesity, adipocyte death in WAT has been shown to promote the accumulation of adipose tissue macrophages (ATM) in crown-like structures (*19*) and triggers their pro-inflammatory phenotype (*20*), which in turn contributes to obesity-associated comorbidities such as type 2 diabetes (*19*, *21–23*). Of note, ATM likely play a dual role in obesity with many macrophages adapting a lipid-associated macrophage (LAM) phenotype due to their lipid-handling capacity. This macrophage subset contributes to metabolic homeostasis (*24*, *25*). Interestingly, LAMs also emerge in visceral WAT during intermittent fasting, as a response to adipocytes undergoing p53-dependent apoptosis, hence suggesting substantial cell death also during adipose tissue catabolic stimulation and shrinkage (*26*).

BAT has been shown to be far less susceptible to macrophage infiltration and inflammatory macrophage induction in general, especially upon high fat diet feeding (*27*). It is thereby unclear, if any of these differences can be attributed to BAT-specific mediators affecting macrophage core functions such as efferocytosis. In BAT, LAM cells regulate brown to white fat conversion by scavenging damaged lipids and mitochondria from metabolically overloaded brown adipocytes via the scavenging receptor CD36 (*28*). Similarly, CD36-mediated scavenging of adipocyte-derived EVs containing damaged mitochondria contributes to efficient thermogenesis (*29*). Additionally, BAT macrophages have been discussed to support thermogenesis directly via the production of norepinephrine (NE), the primary driver of adipose tissue activation (*30*, *31*). The relevance of neuro-immunometabolic signalling has been further confirmed by the role of adipose tissue and sympathetic neuron-associated macrophages (SAMs) in mediating NE bioavailability, therefore regulating energy expenditure and lipolysis (*32*, *33*). Moreover, a subpopulation of CD169^+^ nerve-associated macrophage (NAMs), has been recently shown to protect visceral adipose tissue from age-related dysfunction by preventing inflammation and catecholamine resistance (*34*).

In addition to activating thermogenesis and lipolysis in adipocytes, NE also profoundly affects macrophage biology and function. This includes the promotion of anti-inflammatory polarization (*35*, *36*), control of lipid metabolism (*37*) and migration (*38*), as well as decreasing antimicrobial and antiviral activity (*39*, *40*). Still, whether NE signalling can exert an effect on adipose tissue homeostasis and remodelling by modulating macrophage efferocytosis has not been studied.

Here, in a model of BAT thermogenic remodelling induced by cold exposure, we identified NE as a novel tissue-specific inducer of efferocytosis in macrophages. Activation of the β2-adrenergic receptor on macrophages resulted in increased efferocytosis and induction of the receptor tyrosine kinases *Axl* and *Mertk,* key efferocytic receptors and hence important homeostatic regulators, broadly expressed by macrophages in many tissues (*41*). Furthermore, we demonstrate that mice lacking the expression of the efferocytic receptors *Axl* and *Mertk* in macrophages exhibit impaired thermogenic and lipolytic activation of interscapular BAT (iBAT) and inguinal WAT (ingWAT) in response to cold challenge after high fat diet feeding. Overall, the presented results add to our understanding of neuro-immune crosstalk in the resolution of inflammation, and establish AXL/MERTK-dependent efferocytosis as a factor contributing to adipose tissue remodelling and energy homeostasis.

## RESULTS

### Thermogenic activation stimulates efferocytosis in a lipid-scavenging adipose tissue macrophages subpopulation

Lipid-associated macrophages are described as key responders of adipose tissue metabolic stress and modulators of adipose tissue function tissue function (*24*, *26*, *28*). However, the role of lipid handling by macrophages during brown adipose tissue (BAT) thermogenic activation and remodelling has not yet been fully dissected. To investigate the lipid-handling capacity of BAT macrophages, we first characterized the lipidomic signature of CD11b^+^ cells isolated from the BAT between cold exposed and room temperature housed mice. CD11b⁺ cells isolated from the BAT of cold-exposed animals appeared to accumulate triacylglycerides (TGs) and were characterized by significantly higher levels of phospholipids, particularly phosphatidylethanolamine (PE) species (**Fig. 1A**). To understand origins and consequences of lipid accumulation within BAT macrophages, we reanalysed a previously published ScSeq dataset of murine BAT under conditions of thermoneutral and cold exposure (*42*). Among the 9 clusters identified, the analysis revealed a subcluster of BAT macrophages, cluster 2, that was highly enriched in transcripts related to lipid-handling (**Fig. 1B, C, Supplementary Table 1**), with further enrichment induced by cold exposure, particularly when compared to the thermoneutral condition where BAT activity is minimal (**Fig. 1D**). Flow cytometry analysis performed in cold-exposed BAT, confirmed the presence of a CD11b^+^F4/80^+^LY6C^low^ macrophage population expressing the class B scavenger receptor CD36, CD9, and TREM2 (fig. **S1A**), previously identified as markers of lipid-handling macrophages (*24*, *28*).

**Fig. 1.**
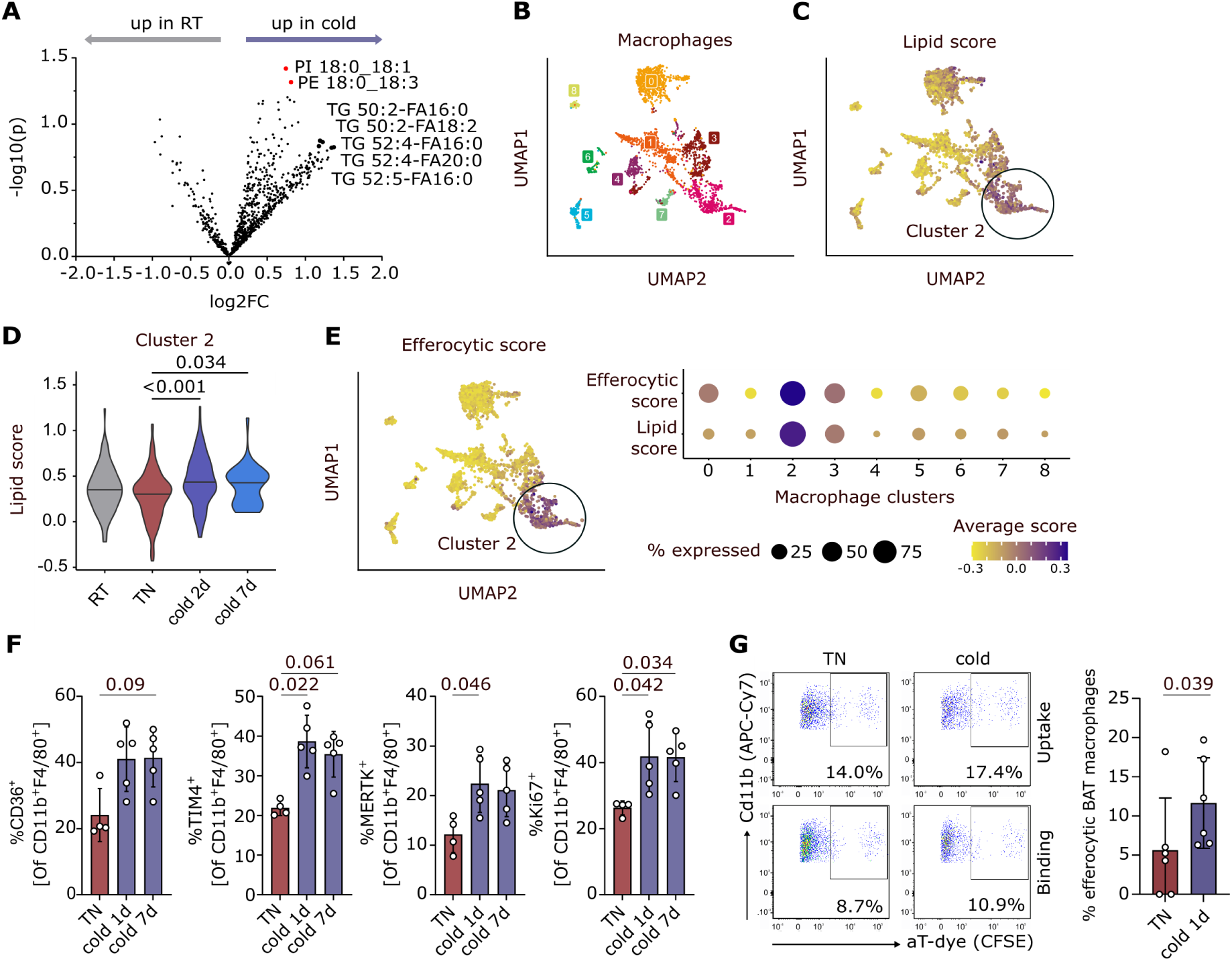
Cold induced-lipid handling macrophages in BAT have increased efferocytic capacity upon cold exposure. (**A**) Volcano plot of lipid species concentration in CD11b^+^ cells from interscapular brown adipose tissue (iBAT), comparing mice kept at room temperature (RT) and mice exposed to cold (6°C) for 1 day. Dots annotate individual lipid species. Significantly changed lipid species are marked in red. n=4. unpaired ttests; (**B**) UMAP reporting macrophage clusters identified in single-cell sequencing data of murine BAT by Shamsi et al. (*42*) (GSE160585) with lipid score overlayed in (**C**); (**D**) Violin plots reporting lipid score of macrophage cluster 2 across temperature conditions. Wilcoxon test, fdr-corrected; (**E**) (left) UMAP reporting efferocytic score among macrophages and (right) dot plot reporting efferocytic and lipid score in the different macrophage clusters as reported in (B). (**F**) Efferocytic receptor expression in CD45^+^CD11b^+^F4/80^+^ iBAT macrophages isolated from mice kept at thermoneutral conditions (30°C) or exposed to cold (6°C) for 1 (1d) or 7 days (7d), as assessed by flow cytometry. Each dot corresponds to one mouse. n=5. Mean±SDs. Kruskal-Wallis tests (p=0.046, p=0.008, 0.027 and 0.009 respectively) followed by Dunn’s multiple comparison tests; (**G**) Representative dot plots and pooled results of *ex vivo* efferocytosis assays in iBAT CD11b^+^ cells isolated from mice kept at thermoneutral conditions (TN) or exposed to cold for 1 day (1d). Each dot represents a pool of 3 mice. n=6. Mean±SD. Mann-Whitney test.

The lipid handling identity of this macrophage subcluster might partially result from the uptake of (phospho)lipids from the extracellular space or from membrane components of dying cells engulfed by macrophages via efferocytosis, as previously described (*43*). Supporting this notion, a high number of efferocytosis-associated transcripts was exclusively detected in the subcluster of macrophages enriched in lipid-handling genes (**Fig. 1E**). Overall, these results suggest that thermogenic activation promotes lipid accumulation in a unique subpopulation of BAT macrophages with efferocytic features.

To evaluate the features of those BAT macrophages *in vivo*, we analyzed by flow cytometry the levels of efferocytic receptors such CD36, TIM4 and the receptor tyrosine kinase MERTK. We observed higher efferocytic receptor expression in interscapular (iBAT) macrophages upon cold exposure (6°C) for 1 day or 7 days as well as higher proliferative capacity as detected by Ki67 labelling (**Fig. 1F**, fig. **S1B, C**). This suggests that efferocytosis-induced proliferation, a process aiming at expanding the pool of resolving macrophages (*44*), might be occurring in BAT during thermogenic activation.

To further prove whether iBAT macrophages retrieved from cold exposed mice have a higher ability to perform efferocytosis, we isolated CD11b^+^ cells via magnet-cell-sorting and performed an efferocytosis assay *ex vivo* by providing exogenous apoptotic cells. Importantly, the frequency of CD11b⁺F4/80⁺ macrophages engulfing pre-labeled apoptotic targets was higher in cells isolated from the iBAT of cold-housed mice compared to controls kept at thermoneutrality (**Fig. 1G**, fig. **S2A**). Although less pronounced, higher efferocytic capacity was observed in macrophages from other adipose tissue depots (inguinal WAT and gonadal WAT) isolated from cold exposed mice (fig. **S2B**). These findings confirm that thermogenic activation, in addition to enhancing lipid handling in a population of BAT efferocytic macrophages, also increases the actual capacity of iBAT macrophages to engulf dying cells.

### Accumulation of dying cells and IL-4 characterizes BAT upon thermogenic activation

To unravel the potential reasons for triggering efferocytosis upon thermogenic activation, we analyzed markers of cellular stress and apoptosis in murine BAT. Bulk mRNA sequencing of stromal vascular cells (SVF) and mature adipocytes (mAdipocytes) revealed alterations in genes involved in the regulation of apoptosis, such as the pro-apoptotic gene *Bax* induced upon cold exposure (**Fig. 2A**). Higher *Bax* levels were further confirmed by qPCR and Western blot (**Fig. 2B**, fig. **S3A**). Similarly, the re-localization of cytochrome C from the mitochondria to the cytosol, a hallmark of apoptosis (*45*), was observed in iBAT upon cold exposure, via compartment-specific proteomic analysis (**Fig. 2C**). Together, these data suggest that thermogenic activation acts as a trigger of cell death within BAT, potentially requiring increased efferocytic activity.

**Fig. 2.**
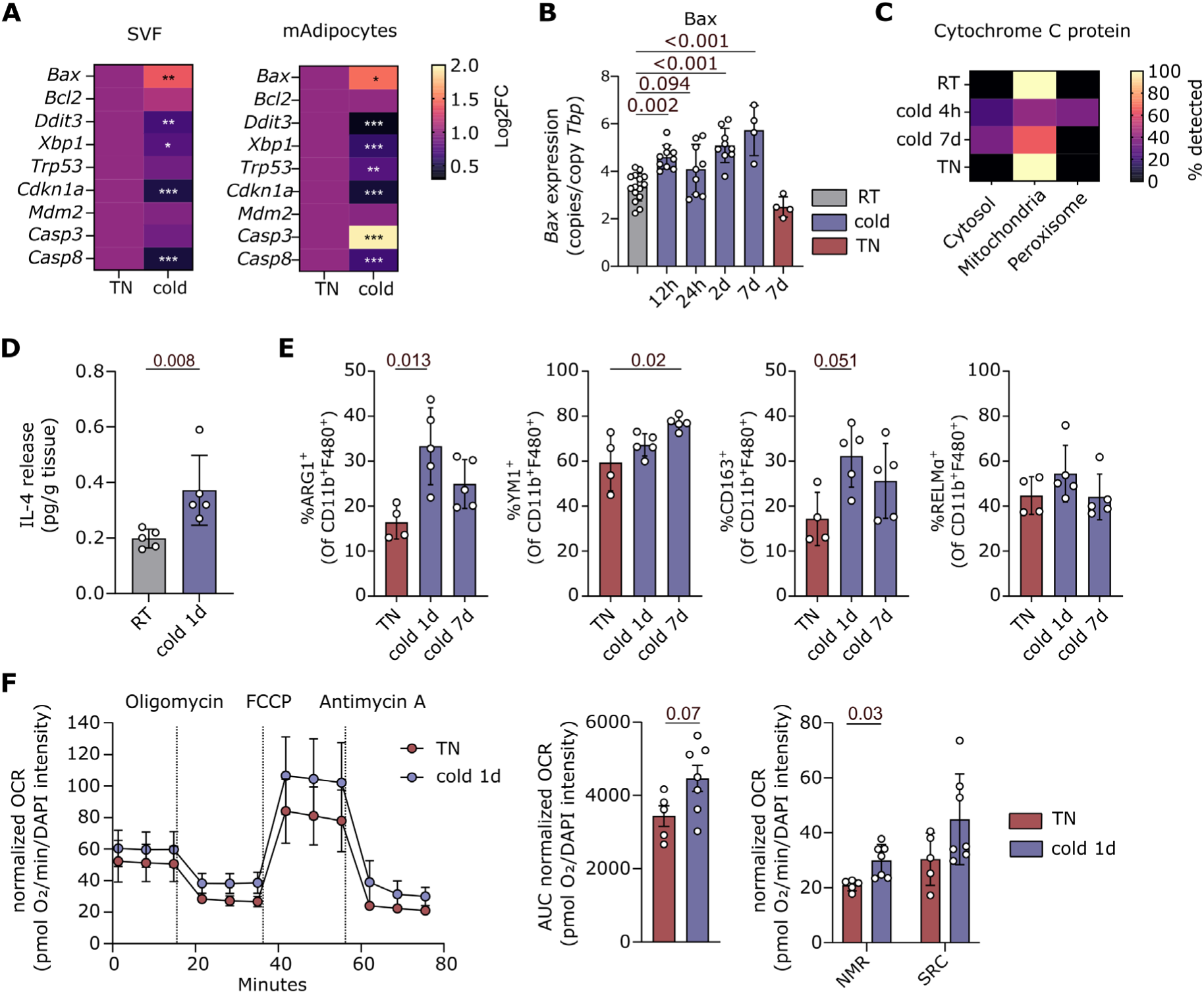
Dying cells and IL-4 accumulate in BAT upon cold exposure. (**A**) mRNA levels of selected apoptosis-related genes in mature adipocyte (mAdipocyte) or stromal vascular fraction (SVF) isolated from interscapular BAT (iBAT) of mice either kept at thermoneutral (TN, 30°C) or exposed to 2 days of cold (6°C) after two weeks of acclimatization to thermoneutral (TN, 30°C) housing conditions. mRNA levels were assessed by bulk mRNA-Sequencing and are displayed as a heatmaps. n=3. *padj ≤ 0.05; **padj ≤ 0.01; ***padj ≤ 0.001; (**B**) *Bax* mRNA expression as assessed by RT-qPCR in whole iBAT of mice either exposed to room temperature (RT), or cold for 12h, 24h, 2, and 7 days (d) or TN for 7 days (d). n=4-18. Kruskal-Wallis test (p<0.0001) followed by Dunn’s multiple comparison test; (**C**) Heatmap indicates distribution of proteins across compartments as predicted by organelle proteomics analysis based on neuronal network based classification. n=1, resulting from a pool of 5 mice per group; (**D**) IL-4 levels in supernatants of iBAT explants, in mice kept at RT or cold conditions for 1 day, as assessed by LEGENDplex analysis. n=5. Mann-Whitney test; (**E**) Flow cytometric analysis of CD45^+^CD11b^+^F4/80^+^ iBAT macrophage phenotype in mice kept at TN conditions or exposed to cold for 1 day or 7 days. n=5. Kruskal-Wallis test (0.008, 0.008, 0.046, 0.254) followed by Dunn’s multiple comparison test; (**F**) Mitochondrial stress test, measured via Seahorse, in iBAT CD11b^+^ cells of isolated from mice kept at TN conditions or exposed to cold for 1 day. n=6/7. Mann-Whitney test. For seahorse analysis 6 mice were pooled to obtain one datapoint. One outlier was detected (ROUT method, Q=1%) and was excluded from analysis. OCR=oxygen consumption rate, AUC=area under the cuve, NMR=non-mitochondrial respiration. SRC=spare respiratory capacity; All data are presented as Mean±SD and each dot corresponds to one mouse, unless stated otherwise.

Given the importance of the coincident detection of dying cells and anti-inflammatory cytokines, such as IL-4, in shaping the function of efferocytic macrophages (*46*), we next analyzed IL-4 levels in BAT explants from mice kept at thermoneutrality or exposed to cold. In our experimental settings, thermogenic activation was associated with IL-4 release (**Fig. 2D**), as previously described (*47*, *48*), along with a tendency towards lower secretion of pro-inflammatory cytokines such as TNFα and IL-6 (fig. **S3B**). In line, CD11b^+^F4/80^+^ macrophages isolated from cold-exposed mice exhibited a stronger anti-inflammatory profile compared to their control counterparts, as reflected by their elevated expression of ARG1, YM1 and CD163, and a tendency towards increased RELMα expression (**Fig. 2E**, **fig S1B** and **S3D**). Functionally, higher respiratory rates were observed in CD11b⁺ cells isolated from the iBAT of cold-exposed mice compared to macrophages isolated from mice kept at thermoneutrality. This was detected as a trend towards higher area under the curve values (AUC) and significantly higher non-mitochondrial respiration during Seahorse analysis (**Fig. 2F**).

Altogether, these data indicate that the increased accumulation of dying cells and IL-4, triggered by thermogenic activation of the BAT, is associated with macrophage phenotypical and cellular reprogramming, as previously described for efferocytic macrophages in other IL-4-dominated environments (*46*, *49*, *50*).

### Norepinephrine is a novel trigger of efferocytosis in macrophages

Tissue-specific signals are critical mediators of macrophage function, driving macrophage heterogeneity in various experimental settings. Examples include adiponectin, released by adipocytes (*17*, *51*) and bile acids in the liver (*52*), both of which have been reported to drive an anti-inflammatory response in macrophages and promote the resolution of inflammation. To identify adipose tissue-specific factors, which may promote efferocytic capacity in BAT macrophages upon cold challenge, we employed an *in vitro* experimental setting in which bone marrow-derived macrophages (BMDMs) were stimulated with conditioned media from primary murine brown adipocytes, which were activated with norepinephrine (NE). NE, normally released by sympathetic neurons, is a key mediator of BAT thermogenic activation (*53*). Treatment of BMDMs with NE-conditioned adipocyte supernatant resulted in higher amounts of triglycerides in macrophages, compared to macrophages treated with conditioned medium from unstimulated adipocytes, as shown by bodipy staining and by lipidomic analysis respectively (**Fig 3A, B**). Critically, NE-conditioned adipocyte supernatant also significantly induced efferocytosis in BMDMs compared to conditioned medium from unstimulated adipocytes (**Fig. 3C**, fig. **S4A**).

**Fig. 3.**
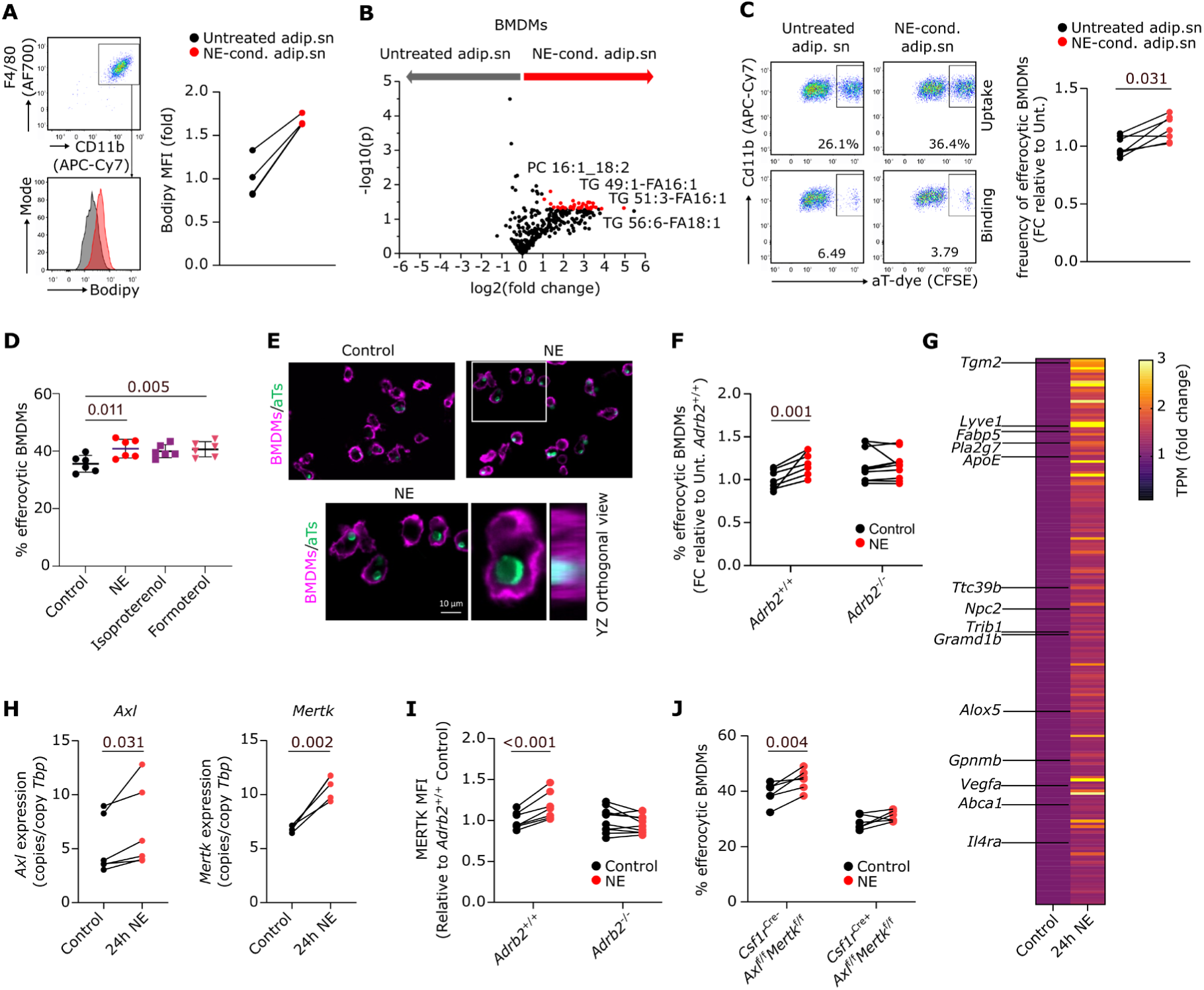
Norepinephrine profoundly shapes macrophage efferocytic function. (**A**) Bodipy staining of CD11b^+^F480^+^ bone marrow-derived macrophages (BMDMs) incubated for 24h with conditioned supernatant (sn) from either untreated brown adipocytes or brown adipocytes stimulated with 1 µM norepinephrine (NE) for 24h prior to supernatant collection. Representative histogram and quantification of mean fluorescence intensity (MFI) are reported. n=3; (**B**) Lipid species concentration in BMDMs treated as in (A). Dots represent individual lipid species. Significantly changed lipid species are marked in red. n=3. unpaired ttest; (**C**) Representative dot plot and pooled results reporting efferocytic capacity of BMDMs treated as in (A). n=7. Wilcoxon test. Efferocytosis in individual samples was normalized to the efferocytosis rate in the untreated adipocyte supernatant condition (**D**). Comparison of efferocytosis capacity between vehicle treated BMDMs and BMDMs treated with 1 µM NE, 1 µM isoprotenerol, or 1 µM formoterol for 24h. n=6. Mean±SD. Friedman test (p=0.002) followed by Dunn’s multiple comparison test. (**E**) Microscope images of BMDMs (magenta) after efferocytosis of apoptotic thymocytes (green) in absence of NE (control) and presence of NE. Single BMDM containing a single apoptotic thymocyte is shown with the YZ orthogonal view. Scale bar: 10 µm; (**F**) Pooled results reporting efferocytic capacity of *Adrb2^+/+^*and *Adrb2^−/−^* BMDMs treated with 1 µM NE for 24h. n=6. Repeated Measures two-way ANOVA followed by Šidák’s multiple comparison test. Efferocytosis in individual samples was normalized to the efferocytosis rate in the control condition for *Adrb2*^+/+^ BMDMs. (**G**) Heatmap reporting the 200 most significantly upregulated genes in BMDMs after 24h treatment with 1 µM NE. n=3; (**H**) Normalized gene expression of *Axl* and *Mertk* in BMDMs treated with 1 µM NE for 24h. n=6/5. Mann-Whitney test; MFI for MERTK in *Adrb2*^+/+^ and *Adrb2*^−/−^ BMDMs (**I**) and Efferocytosis capacity of *Csf1r^Cre-^Axl^f/f^Mertk^f/f^*and *Csf1r^Cre+^Axl^f/f^Mertk^f/f^* BMDMs (**J**) after treatment with 1 µM NE for 24h. n=7/10 and n=6 respectively. Repeated Measures two-way ANOVA followed by Šidák’s multiple comparison test; Each dot represents one biological replicate and paired replicates are connected with a black line.

Based on this observation, we hypothesized that NE in the NE-conditioned adipocyte supernatant might be sufficient to induce efferocytosis in macrophages. To test this, we treated BMDMs directly with NE or other β2-adrenergic agonists. Indeed, direct NE stimulation of BMDMs was sufficient to significantly induce efferocytosis in BMDMs (**Fig. 3D**), suggesting that NE might serve as a physiological trigger of cold-induced efferocytosis *in vivo.* Moreover, efferocytosis was also induced by stimulation with isoprotenerol (a non-selective β-adrenergic agonist) or formoterol (a β2-selective agonist), establishing β2-adrenergic signaling in general, rather than NE specifically, as a novel inducer of efferocytosis in macrophages (**Fig. 3D**). To corroborate these observations and confirm the uptake of dying cells, we used microscopy. We observed that dying cells were fully engulfed after NE treatment as shown in 3D and exemplified with an orthogonal view (**Fig. 3E**). In line with the data on β2-adrenergic stimulation, NE failed to promote efferocytosis in BMDMs with genetic ablation of the β2-adrenergic receptor (**Fig. 3F**). This identifies the β2-adrenergic receptor signaling as both necessary and sufficient for the effect of NE on efferocytosis. In line with this, analysis of sc-RNAseq data from BAT macrophages isolated from cold-exposed mice (*42*) confirmed the enrichment of the β2-adrenergic receptors expression in the subcluster of efferocytic/lipid-handling macrophages, as reported in Fig. 1. This cluster was characterized by the expression of *Maoe*, suggesting features of sympathetic neuron-associated macrophages, which have previously been described for their role in NE scavenging and NE-mediated regulation of thermogenesis (fig. **S3B**) (*33*).

To investigate the molecular mechanism mediating NE-induced efferocytosis, we performed bulk mRNA sequencing of BMDMs following NE stimulation. BMDMs treated with NE for 24 hours clustered separately from untreated BMDMs, indicating a distinct transcriptional program (fig. **S4C**). NE strongly modulated macrophages response with 558 genes being significantly up and 324 genes being significantly downregulated (**Supplementary Tables 2**). The IL-4 receptor (*IL4ra*), genes involved in angiogenesis (e.g *Vegfa*), extracellular matrix binding (e.g. *Lyve1*), as well as lipid processing genes (*Alox5, Pla2g7, Gpnm*) were among the most significantly upregulated ones. Interestingly, LXR target genes and cholesterol metabolism genes (*Tgm2, Fabp5, ApoE, Abca1, Ttc39b, Npc2, Trib1, Gramd1b*) were the 200 most significantly upregulated genes after 24 hours of NE treatment (**Fig. 3G**), suggesting that the response to NE drives alterations in macrophage polarization status, which are associated with the regulation of both lipid metabolism and efferocytic function.

Importantly, the higher efferocytic capacity observed upon treatment of BMDMs with NE was associated with increased mRNA expression of the efferocytic receptors *Axl* and *Mertk*, which together with Tyro3, belong to the family of receptor tyrosine kinases known as TAM (*54*), as detected by qPCR (**Fig. 3H**). This finding aligns with increased MERTK protein levels in BAT macrophages following cold exposure (Fig. 1E), as well as with elevated *Mertk* mRNA expression in CD11b⁺ BAT cells isolated from cold-exposed mice (fig. **S4D**). Additionally, cold exposure induced *Abca1* mRNA expression, a key LXR target (*55*) in BAT CD11b^+^ cells (fig. **S4D**), suggesting that cold-induced *Mertk* expression *in vivo* may indeed be linked to LXR signaling, as previously described in other settings (*14*). MERTK expression was triggered by NE sensing through the β2-adrenergic receptor, as demonstrated by increased expression of MERTK in *Adrb2*^+/+^ but not *Adrb2*^−/−^ mice after 24h of treatment (**Fig. 3I**). To further confirm whether NE regulates macrophage efferocytic capacity through MERTK, we provided apoptotic cells to BMDMs differentiated from either *Csf1r^Cre-^Axl^f/f^Mertk^f/f^*or *Csf1C^Cre+^Axl^f/f^Mertk^f/f^* mice (*56*, *57*), in which *Csf1r^Cre+^Axl^f/f^ Mertk^f/f^* macrophages lack the expression of AXL and MERTK receptors. In these macrophages, NE failed to enhance their efferocytic capacity (**Fig. 3J**).

Altogether, these data indicate that NE, upon being sensed by the β2-adrenergic receptor, acts as a trigger of macrophage efferocytosis through engagement of the TAM receptors MERTK and AXL.

### NE triggers monocyte/macrophage efferocytic profile *in vivo*

Building on the *in vitro* effects of NE on efferocytic macrophages, we next investigated whether NE could serve as a physiological trigger of efferocytosis *in vivo*. To this end, wild type (WT) mice were injected with NE and BAT macrophages were analyzed for their *ex vivo* capacity to engulf dying cells. Indeed, NE treatment resulted in a trend towards higher efferocytosis in CD11b^+^F4/80^+^ BAT macrophages (**Fig. 4A**, fig. **S4E**), with stronger effects on a subset of CD11b^+^F4/80^+^CX3CR1^+^LY6C^low^ BAT macrophages (fig. **S4F**). Moreover, NE injection resulted in elevated expression of efferocytic receptors, most notably AXL and MERTK, but also TIM4 and CD36, especially within a population of non-classical monocytes (**Fig. 4B**, fig. **S4G, H**). Taken together, these results indicate that NE can shape the efferocytic signature of circulating monocytes and, through its sensing in BAT, renders BAT macrophages more prone to engulf dying cells.

**Fig. 4.**
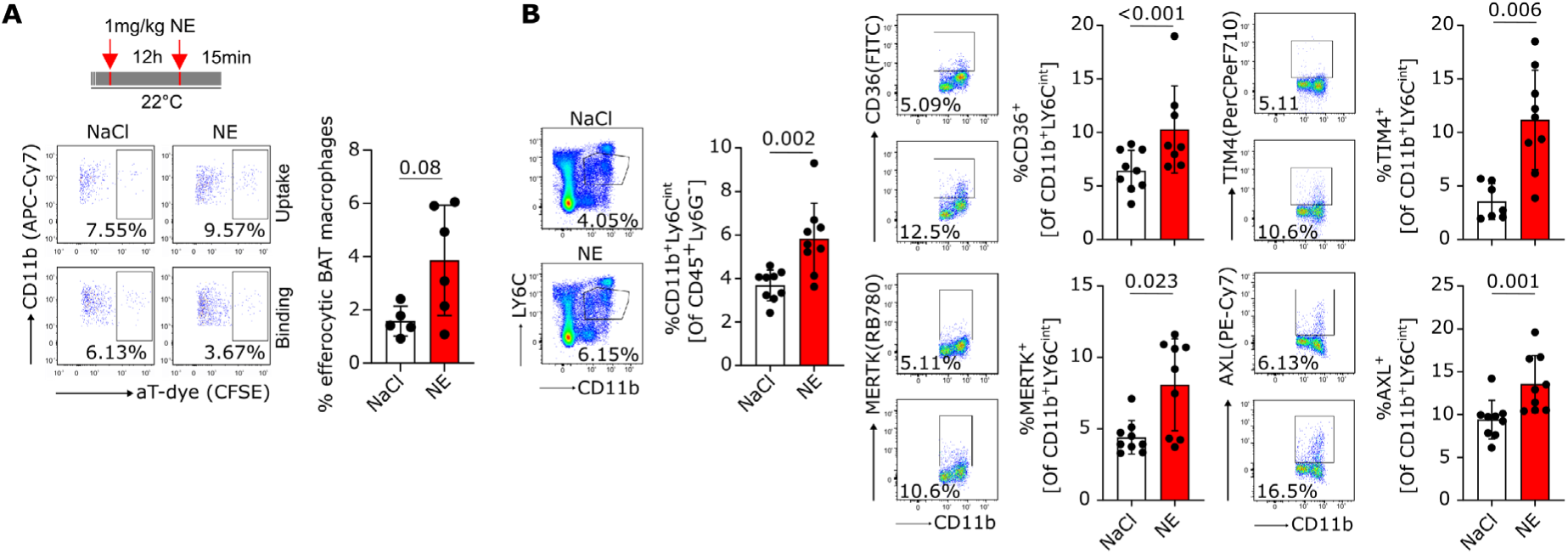
NE directly enhances efferocytosis receptor expression in vivo. (**A**) Experimental layout (upper panel). WT mice were injected subcutaneously with natrium-chlorid (NaCl) or norepinephrine (NE) at 1 mg/kg bodyweight twice, 12 h and 15 minutes before analysis. Pooled results (bottom panel) reporting *ex vivo* efferocytosis assays of Live^+^CD45^+^CD11b^+^F480^+^ interscapular BAT (iBAT) macrophages isolated from mice treated with NaCl or NE. Each dot represents a pool of 3 mice. n=6. Mean±SD. Mann-Whitney test. One outlier was detected (ROUT method, Q=1%) and was excluded from analysis; (**B**) Efferocytic receptor expression in CD45^+^LY6G^low^CD11b^+^Ly6C^int^ non-classical blood monocytes in mice after injection with NaCl or NE at 1mg/kg bodyweight twice, 12 h and 15 minutes before analysis as in (A). n=7-9. Mean±SD. Mann-Whitney tests; Outlier tests were performed for each epitope separated using ROUT method (Q=1%) and excluded from the corresponding graph.

### AXL/MERTK-dependent efferocytosis regulates adipocyte thermogenic and lipolytic functions

To further investigate the effects of NE-induced efferocytosis via engagement of *Axl* and *Mertk* on adipose tissues and systemic metabolism, we housed *Csf1r*^Cre-^*Axl*^f/f^*Mertk*^f/f^ and *Csf1r*^Cre+^*Axl*^f/f^*Mertk*^f/f^ at thermoneutral conditions (30°C) and fed the mice a high-fat diet (HFD). Applying these “physiologically humanized” conditions, result in the presence of enlarged unilocular adipocytes and moderate immune cell infiltration, including macrophages, into BAT (*58*, *59*). Of note, these morphologic and concomitant transcriptional changes are considered to phenocopy human BAT. We subsequently acclimated the mice to two cycles of cold exposure, to elicit a recurring and robust influx of NE into the BAT. Phenotypically, BAT macrophages from *Csf1r*^Cre+^*Axl*^f/f^*Mertk*^f/f^ were similar to BAT macrophages from control *Csf1r*^Cre-^*Axl*^f/f^*Mertk^f^*^/f^ mice (fig. **S5A**-**C**) and cytokine secretion of BAT explants as well as pro-inflammatory gene expression remained unchanged (fig. **S5D**). However, transcription of genes involved in adaptive thermogenesis and lipid handling such as *Ucp1*, *Prdm16* and *Lpl* was lower in BAT from *Csf1r*^Cre+^*Axl*^f/f^*Mertk*^f/f^ mice compared to controls, suggesting impaired thermogenic capacity in a setting with limited efferocytosis (**Fig. 5A**). Of note, these effects were not due to altered iBAT innervation in the two mouse strains, as confirmed by similar levels of tyrosine hydroxylase (TH), the main NE synthesizing hormone, assessed by Western blotting (fig. **S6A**). Nonetheless, the effects observed on thermogenic mRNA expression did not result in lower UCP1 protein levels in BAT and did not strongly affect iBAT morphology (fig. **S6B** and **C**).

**Fig. 5.**
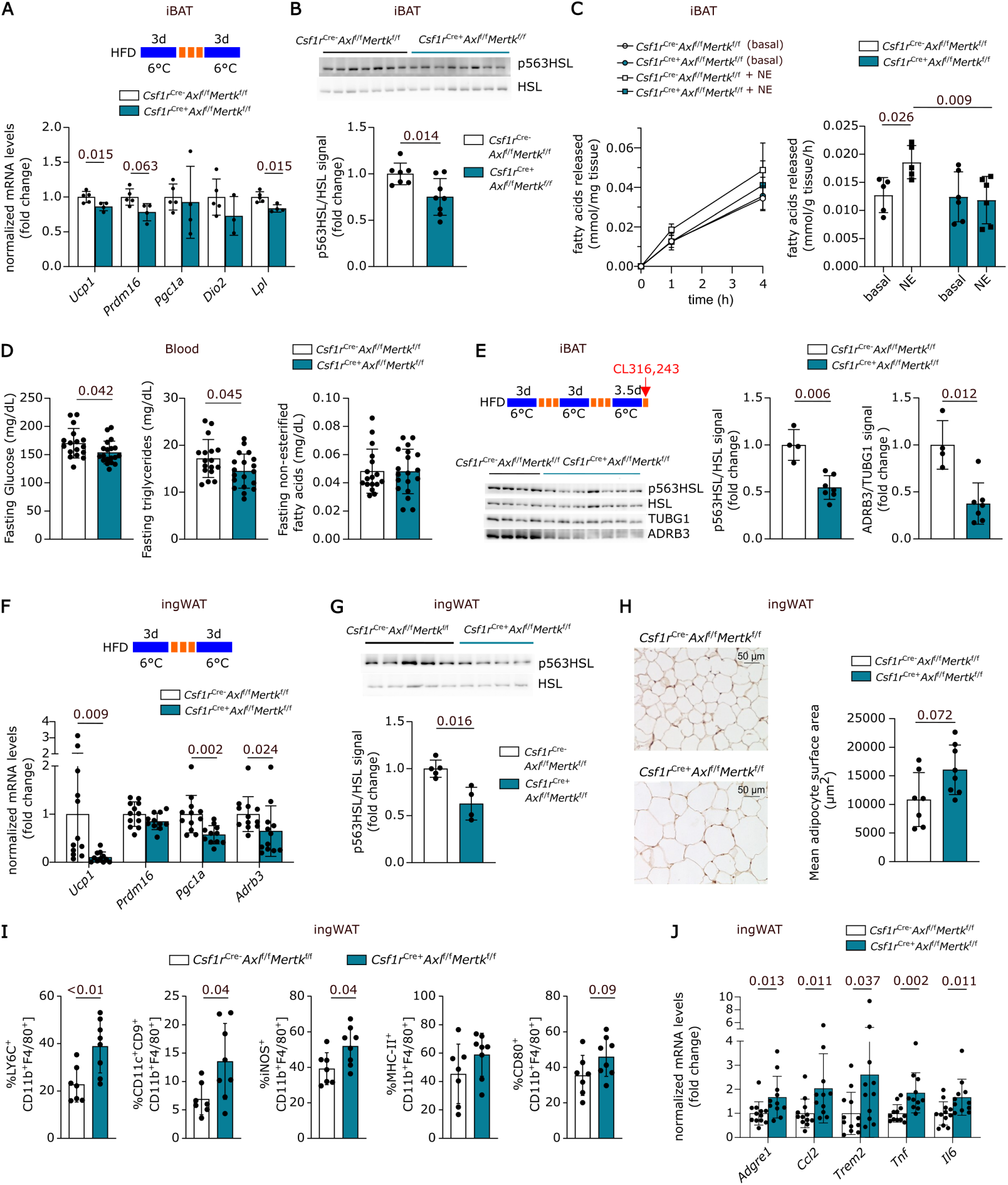
AXL/MERTK-dependent efferocytosis controls adipose tissue activation and inflammation after high fat diet and repeated cold challenge. (**A**) Experimental layout (upper panel) and expression of thermogenic genes in interscapular BAT (iBAT) of *Csf1r*^Cre-^*Axl*^f/f^*Mertk*^f/f^ and *Csf1r*^Cre+^*Axl*^f/f^*Mertk*^f/f^ mice after high fat diet (HFD) and repeated cold exposure (bottom). n=5/4. Mann-Whitney tests. Results were normalized to the mean expression in *Csf1r*^Cre-^*Axl*^f/f^*Mertk*^f/f^ mice. (**B**) Phosphorylated HSL (p563HSL) to total HSL ratio in iBAT of *Csf1r*^Cre-^*Axl*^f/f^ *Mertk*^f/f^ and *Csf1r*^Cre+^*Axl*^f/f^*Mertk*^f/f^ mice treated as in (A). n=7/8. Mann-Whitney test. (**C**) Secretion of non-esterified fatty acids (NEFAs) by iBAT explants isolated from *Csf1r*^Cre-^*Axl*^f/f^*Mertk*^f/f^ and *Csf1r*^Cre+^*Axl*^f/f^*Mertk*^f/f^ mice after HFD and repeated cold exposure after 1h and 4h incubation with and without stimulation with 1 µM NE. n=5/6. 2-way ANOVA followed by uncorrected Fisher’s LSD test; (**D**) Fasting glucose, triglycerides and NEFAs in plasma of *Csf1r*^Cre-^*Axl*^f/f^*Mertk*^f/f^ and *Csf1r*^Cre+^*Axl*^f/f^*Mertk*^f/f^ mice treated as in (A). n=17-19. unpaired ttests. (**E**) Experimental layout (upper panel) and p563HSL to total HSL ratio and ADRβ3 to gamma tubulin (TUBG1) ratio in iBAT of *Csf1r*^Cre-^*Axl*^f/f^*Mertk*^f/f^ and *Csf1r*^Cre+^*Axl*^f/f^*Mertk*^f/f^ mice (bottom panel). iBATs were collected after HFD feeding, repeated cold exposure and a final injection of CL316,243 at 1mg/kg bodyweight five h before organ harvest. n=4-7. Mann-Whitney tests. (**F**) Experimental layout (upper panel) and expression of thermogenic genes in ingWAT of *Csf1r*^Cre-^*Axl*^f/f^*Mertk*^f/f^ and *Csf1r*^Cre+^*Axl*^f/f^*Mertk*^f/f^ mice after HFD treated as in (A) (bottom panel). n=11-12. Mann-Whitney tests. (**G**) p563HSL to total HSL ratio in ingWAT of *Csf1r*^Cre-^*Axl*^f/f^ *Mertk*^f/f^ and *Csf1r*^Cre+^*Axl*^f/f^ *Mertk*^f/f^ mice after HFD treated as in (A). n=4-5. Mann-Whitney test. (**H**) Hematoxylin and UCP1 staining to quantify mean lipid droplet size in ingWAT of *Csf1r*^Cre-^*Axl*^f/f^*Mertk*^f/f^ and *Csf1r*^Cre+^*Axl*^f/f^*Mertk*^f/f^ mice treated as in (A). n=7/8. Mann-Whitney test. (**I**) Flow cytometric analysis reporting the inflammatory status of Live^+^CD45^+^LY6G^low^CD11b^+^F4/80^+^ ingWAT macrophages from *Csf1r*^Cre-^*Axl*^f/f^*Mertk*^f/f^ and *Csf1r*^Cre+^*Axl*^f/f^*Mertk*^f/f^ mice treated as in (A). n=7-8. Mann-Whitney tests. (**J**) Expression of inflammatory genes in ingWAT of *Csf1r*^Cre-^*Axl*^f/f^*Mertk*^f/f^ and *Csf1r*^Cre+^*Axl*^f/f^*Mertk*^f/f^ mice treated as in (A). n=11-12. Mann-Whitney tests. In this Figure, each dot corresponds to one mouse and all data are displayed as Mean±SD; In (B), (E-G), (J), results were normalized to the mean expression in *Csf1r*^Cre-^*Axl*^f/f^*Mertk*^f/f^ mice.

In response to beta-adrenergic stimulation, as in the case of NE influx upon cold exposure, adipose tissues not only enhance a thermogenic program, but also liberate free fatty acids from TAG stores within lipid droplets, via phosphorylation of hormone-sensitive lipase (HSL) (*60*). Of note, we observed significantly lower levels of HSL phosphorylation within iBAT of mice lacking *Axl* and *Mertk* expression in macrophages (*Csf1r*^Cre+^*Axl*^f/f^*Mertk*^f/f^), suggesting diminished sympathetic activation of adipocytes (**Fig. 5B**). Importantly, the reduction of HSL phosphorylation *in vivo* was accompanied by a lower ability of BAT explants from *Csf1r*^Cre+^*Axl*^f/f^ *Mertk*^f/f^ mice to perform NE-induced lipolysis *ex vivo,* indicating catecholamine resistance (**Fig. 5C**). Those effects on lipolysis were associated to significantly lower fasting glucose and triglyceride levels in the blood of *Csf1r*^Cre+^*Axl*^f/f^ *Mertk*^f/f^ mice but no changes in non-esterified fatty acids (**Fig 5D**).

Together, these results suggest that the impairment in macrophage efferocytosis via *Axl* and *Mertk* renders adipocytes unable to properly react to the adrenergic stimulus, subsequently affecting thermogenic gene expression and lipolysis. To confirm this notion *in vivo*, *Csf1r*^Cre+^*Axl*^f/f^*Mertk*^f/f^ that have undergone HFD at thermoneutral conditions and were subjected to repeated cold exposure, were treated with the β3 adrenergic agonist CL316,243 to maximally activate adipose tissue in an innervation-independent manner. Also in this setting, absence of macrophage *Axl* and *Mertk* resulted in lower phosphorylation of HSL and additionally lower amounts of ADRB3 (β3-adrenergic receptor) protein levels in iBAT (**Fig. 5E**), as well a trend towards reduced phosphorylation of PKA substrates (fig. **S6D**) in BAT of *Csf1r*^Cre+^*Axl*^f/f^*Mertk*^f/f^ mice compared to their Csf1r^Cre-^ counterparts. This is in line with a reduced ability of adipocytes in the *Csf1r*^Cre+^*Axl*^f/f^*Mertk*^f/f^ to react to adrenergic stimuli. However, whole body energy expenditure during cold exposure or after CL injection remained unaffected (fig. **S6E**).

Similarly to what was observed in BAT, lower expression of key thermogenic markers was detected in inguinal WAT (ingWAT) of mice lacking *Axl* and *Mertk* expression in macrophages. This was reflected by lower mRNA levels of key thermogenic markers, including *Ucp1*, *Prdm16*, and *Pgc-1α*, as well as a lower expression of *Adrb3* (*β3*-adrenergic receptor) and diminished HSL phosphorylation in *Csf1r*^Cre+^*Axl*^f/f^*Mertk*^f/f^ compared to the *Csf1r*^Cre-^*Axl*^f/f^*Mertk*^f/f^ counterparts (**Fig. 5F, G**). Additionally, *Axl* and *Mertk* genetic deletion in macrophages resulted in a trend towards larger ingWAT adipocytes (**Fig. 5H**) after repeated cold exposure and significantly reduced levels of pHSL in response to CL316,243 in ingWAT (fig. **S6F**), consistent with what was observed in in iBAT.

Similar to findings in iBAT, TH levels of *Csf1r*^Cre-^*Axl*^f/f^*Mertk*^f/f^ and *Csf1r*^Cre+^*Axl*^f/f^*Mertk*^f/f^ mice remained unchanged in ingWAT (fig. **S6G**). However, MAC2 (Galectin-3) levels were significantly higher in ingWAT of *Csf1r*^Cre+^ mice (fig. **S6G**), indicating substantial macrophage infiltration into the tissue. Alterations in thermogenic and lipolytic activity was also associated with a higher presence of monocyte-derived Ly6C^+^ macrophages and an accumulation of CD11c^+^CD9^+^ macrophages that can usually be found in crown-like structures surrounding dead adipocytes (*61*) in the ingWAT of *Csf1r*^Cre+^*Axl*^f/f^*Mertk*^f/f^ mice (**Fig. 5I**). Macrophage infiltration was also confirmed by increased mRNA levels of *Adgre1* (encoding for the F4/80 antigen), *Ccl2*, a monocyte-attracting chemokine, and *Trem2* (**Fig. 5H**). Moreover, macrophages in ingWAT exhibited a more pronounced pro-inflammatory status, as indicated by elevated iNOS levels and a trend towards higher expression of antigen-presenting molecules, including MHC-II and CD80 (**Fig.5I**, fig **S7A** and **B**). This more pro-inflammatory macrophage profile was also reflected by significantly higher mRNA levels of the pro-inflammatory cytokines *Tnf* and *Il6* in whole ingWAT (**Fig. 5J**).

Collectively, these findings indicate that macrophage efferocytosis mediated by *Axl* and *Mert*k maintains adipocyte thermogenic and lipolytic capacity. The coordination of adipocyte activity by macrophages occurs in response to NE exposure, which, in turn, enhances macrophage efferocytic capacity and prevents the establishment of a pro-inflammatory environment.

## DISCUSSION

Efferocytosis by macrophages plays a critical role in various pathological conditions (*62*) and its modulation can be influenced by diverse spatial and temporal cues (*14*, *17*, *51*, *63*). Our study identifies norepinephrine (NE) signaling as a previously unrecognized and potent trigger of macrophage efferocytosis. Additionally, it establishes efferocytosis as essential for maintaining adipose tissue homeostasis and function during thermogenic remodelling. Specifically, we describe a lipid handling population of macrophages in cold-activated murine BAT, that expresses high levels of efferocytosis-related transcripts. This population also expresses markers typically associated with sympathetic neuron-associated macrophages which have been described to interact with, and catabolize, NE in murine WAT (*33*). Building on this data, our *ex vivo* and *in vitro* analysis support the notion that NE and β2-adrenergic signalling not only act as established regulators of macrophage lipid handling (*37*), but also serve as key triggers of efferocytosis in bone marrow-derived (BMDMs) and BAT macrophages. We therefore propose that NE induced-lipid handling and efferocytosis represent two coordinated pathways that enable macrophages to efficiently clear lipid-rich dying cells.

The anti-inflammatory effects of adrenergic signalling in macrophages have been previously reported (*40*, *64*, *65*): β2-signaling in intestinal muscularis macrophages has been shown to limit neuronal damage following enteric helminth infection (*66*) and genetic ablation of β2-adrenergic receptors in macrophages leads to increased severity of renal ischemia-reperfusion injury (*67*). Leukocyte-expressed β2-adrenergic receptors are also essential for survival after acute myocardial injury (*68*) and interestingly, MERTK-dependent efferocytosis by monocytic myeloid-derived suppressor cells is important for resolving ischemia/reperfusion injury after lung transplantation (*69*). Ischemia thus represents a context in which both macrophage β2-adrenergic signaling and MERTK-dependent efferocytosis have been reported to be involved, suggesting potential interaction between these pathways.

Our proposed axis linking adrenergic signaling to efferocytosis and tissue remodeling may hold evolutionary significance, given the role of sympathetic nervous system signaling in stress responses that prompt physiological adaptation. In adipose tissue in particular, stress signals lead to rapid energy mobilization through lipolysis, heat production and preparation for high metabolic demands. This process results in cellular loss in adipose tissue, requiring efficient clearance of cellular debris. In this scenario, macrophages scavenging lipids are essential to limit the generation of a pro-inflammatory milieu (*70*). NE sensing by adipose tissue macrophages, occurring during cold stress and metabolic challenges, may represent a fast and localized mechanism to aid macrophages in their response to dying cells and to initiate an efferocytosis-dependent tissue-remodeling response. Consistent with this notion, we speculate that impairment in NE-mediated efferocytosis results in the generation of a pro-inflammatory environment in adipose tissue, similar to what has been reported upon depletion of CD169^+^ neuron associated macrophages during aging (*34*). At the same time we propose that locally higher levels of NE in BAT, compared to WAT, may result in more effcient efferocytosis (*71*), which in turn could contribute to the increased resistance to inflammation reported for BAT (*27*).

The data showing increased AXL and MERTK expression in non-classical monocytes following NE injection also supports the potential for systemic effects of adrenergic signaling on macrophages. In line with this, chronic systemic treatment with isoproterenol has been associated with elevated MERTK expression in cardiac macrophages in the context of heart failure (*72*). Notably, systemic administration of non-selective β2 agonists is widely used for cardiovascular effects (*73*), while selective β2 agonists are often administered locally and sometimes systemically to achieve bronchodilation in obstructive lung diseases (*74*). This reveals a potential for repurposing existing drugs to enhance efferocytosis in treating conditions that could benefit from this.

From a mechanistic point of view, we propose that efferocytosis regulation by NE involves LXR signalling as LXR is a key regulator of macrophage efferocytosis (*14*) and expression of several LXR targets was increased following NE treatment of BMDMs and in BAT macrophages after cold exposure. NE is known to modulate lipid metabolism in macrophages (*37*) and LXR is sensitive to changes in the lipid environment, particularly to oxysterols derived from apoptotic cells (*14*). LXR signalling therefore represents a mechanistic link between macrophage lipid handling and efferocytic capacity (*75*). This underscores the potential role of LAMs as a macrophage subset specialized not only in lipid handling but also in efferocytosis. At the same time, this highlights the potential of adipose tissue as a site where integration of adrenergic signaling, lipid handling, and macrophage-mediated efferocytosis supports tissue remodeling.

Although macrophage β2-adrenergic receptor signaling has been previously described to be dispensable for adipose tissue inflammation and function (*76*), our data suggest that specific adaptation to thermoneutral housing as well as HFD feeding prior to cold exposure, are necessary for dissecting the role AXL/MERTK-dependent efferocytosis in maintaining adipose tissue homeostasis.

In this scenario, we propose that the impaired adipose tissue activation observed in *Csf1r*^Cre+^*Axl^f^*^/f^*Mertk*^f/f^ mice is mediated by a pro-inflammatory environment generated as a result of impaired dead cell removal. Given the role of TNFα in suppressing lipolytic activation of adipose tissue by decreasing expression of the adrenergic β3 receptor on adipocytes (*77*), and considering the increased expression of TNF observed in the ingWAT of *Csf1r*^Cre+^*Axl^f^*^/f^*Mertk*^f/f^ mice, it is tempting to speculate that TNFα signalling may contribute to the impairment in adipose tissue activation observed in our experimental setting.

The discovery of AXL/MERTK-dependent efferocytosis as a contributor to adipose tissue homeostasis and activation significantly supports the hypothesis that impaired efferocytosis is a potential driver of adipose tissue inflammation and dysfunction (*19*, *78*). Importantly, our identification of adrenergic signalling as a novel and upstream regulator of AXL and MERTK-dependent efferocytosis in macrophages reveals a previously unrecognized immunometabolic axis. All in all, these findings not only deepen our understanding of how ATMs re-establish adipose tissue homeostasis upon damage, but also highlight adrenergic signalling as a critical modulator of efferocytosis. Increasing efferocytosis by targeting this pathway could help to limit inflammation and may hold therapeutic potential in diseases characterized by pathological accumulation of dead cells, most prominently metabolic syndrome and type 2 diabetes (*19*, *79*, *80*).

## MATERIALS AND METHODS

### Mice

All mouse experiments were approved by the Animal Welfare Officers of University Medical Centre Hamburg-Eppendorf (UKE) and Behörde für Gesundheit und Verbraucherschutz Hamburg. Wild type mice on a C57BL/6 genetic background were housed and bred in the animal facility of the UKE at 22°C under a day-night cycle of 12h and ad libitum access to food and drinking water. Mice with a total deletion of the Adrenergic beta 2 receptor as well as their control littermates (*Adrb2^+/+^* and *Adrb2^−/−^* mice, Adrb2tm1Bkk/J; the Jackson Laboratory, no. 031496) are on a FVB/129 genetic background and were kindly gifted from the lab of J. Keller (UKE) (*81*). Mice with a macrophage specific deletion of *Axl* and *Mertk (Csf1r^Cre-^ Axl^f/f^Mertk^f/f^* or *Csf1r^Cre+^ Axl^f/f^Mertk^f/f^* mice) on a C57BL/6 genetic background were kindly provided by Carla V. Rothlin (Yale University) (*56*, *57*). Mice were housed and bred in the animal facility at the UKE at 22°C under a day-night cycle of 12h and ad libitum access to food and drinking water.

During the experiments, mice were housed individually with wood bedding in a light- and temperature-controlled facility with a 12h/12h dark regime. Unless stated otherwise, mice were housed at 22 °C (RT) and were fed a standard laboratory chow diet ad libitum (19.10% protein, 4% fat, 6% fiber, from Altromin Spezialfutter GmbH & Co. KG, Germany). All experiments were started when mice were between 8 and 12 weeks of age. At the end of the experiments mice were sacrificed by a lethal dose (15μL/g mouse bodyweight) of a mixture containing Ketamin (23mg/mL)/Xylazin (0.2%) in 0.9% NaCl and after a 4h fasting period. Systemic blood was withdrawn by cardiac puncture with syringes containing 5μL 0.5M EDTA (Sigma-Aldrich). Subsequently, animals were perfused with PBS (Thermo Fisher Scientific) and their harvested tissues were immediately either conserved in 3.7% formaldehyde solution, processed to isolate ATMs, or were stored at −80°C for further use.

### Cold challenge

For cold challenges, animals were housed at 6°C for a period of 1-7 days while keeping their respective diets and housing conditions. Corresponding control groups were kept under identical housing conditions, except for the environmental temperature.

### High fat diet (HFD) feeding

Mice were kept under their respective housing conditions with the diet switched to a high fat diet (HFD) containing 35% lard (Bio-Serv, F3282, 20.5% Protein; 36,0% Fat; 35,7% Carbohydrates; 0% Fiber Fat). HFD feeding was started when mice were 8-12 weeks of age and was carried out under thermoneutral temperature housing conditions (30°C). The diet was kept over the whole course of the experiment, including during cold challenges. During the feeding period, food intake was monitored and mice were weighed once a week. At the end of the experiments blood glucose was determined using an Accu-Chek Aviva (Roche) after a regular 4h fasting period.

### Norepinephrine/CL316,243 injections

5µl/g bodyweight of a 200µg/ml norepinephrine (Cayman Chemicals) 0.9% NaCl solution were injected subcutaneously into the interscapular region of mice, for a total dose of 1mg/kg bodyweight. Injections were carried out twice, 12h and 30 minutes before sacrificing the mice. During the experiments, mice were housed at RT under standard housing conditions. To induce adipocyte specific thermogenesis, 10 µl/g bodyweight of a 100 µg/ml CL316,243 (Tocris Bioscience) 0.9% NaCl solution was injected subcutaneously into the interscapular region of mice for a total dose of 1mg/kg bodyweight. The injection was carried out at the indicated timepoints, while mice were kept in metabolic cages at thermoneutral temperature conditions (28°C).

### Flow cytometry

For flow cytometric analysis of adipose tissue macrophages, adipose tissues were minced into a fine paste and digested in a solution containing 10 mg/mL Collagenase Type II (Sigma Aldrich, Germany) and 0.5% BSA in HBSS supplemented with Ca²⁺ and Mg²⁺ for 40 minutes under constant shaking. EDTA was then added to a final concentration of 10 mM, followed by an additional 10-minute incubation at 37°C with shaking. The resulting cell suspension was filtered through a 100 μm nylon mesh. After two washing steps (10 min at 500 × g), the cells were filtered again through a 70 μm filter. A subsequent centrifugation step was performed to separate the floating adipocyte fraction from the stromal vascular cell (SVC) pellet. The floating layer was carefully removed using a Pasteur pipette. The SVC pellet was gently resuspended (without vortexing) in 500 μL of RBC lysis buffer for 3 minutes, followed by the addition of 10 mL of FACS buffer. After one final centrifugation step, the cells were ready for staining. To block unspecific binding Fc block (BioLegend, anti-CD16/CD32 no.101301) diluted 1:10000 in PBS was added for 15 min at 4°C before the extracellular staining. Washing steps with PBS + 2% FBS were repeated three times and performed after each antibody incubation. For each washing step or change of buffer, cells were pelleted at 800 x g for 2 min. For the staining of surface epitopes, cells were incubated with the antibody cocktail for 35 min at 4°C in the dark. If a viability staining was included in the analysis, viability dye (Invitrogen, L34961) was added diluted in PBS before the surface staining. If intracellular epitopes were analyzed, cells were fixed using 2% paraformaldehyde (Electron Microscopy Sciences) for 30 min at 4°C and permeabilised (BD Biosciences Perm/Wash no. 554723) for 15 min at RT before staining. Intracellular antibodies as well as secondary antibodies were diluted in permeabilisation buffer and staining was performed for 40 min at 4°C in the dark. Bodipy staining was carried out after antibody staining and cell fixation using incubation for x minutes with xM bodipy (ref) dissolved in PBS shortly before analysis. Data was acquired either on a FACSymphony™ A3 Cell Analyzer (BDBiosciences) or on a Cytek® Aurora. All data was analyzed using the FlowJo software V10 (Tree Star). Antibodies used for flow cytometry were purchased from commercial sources.

### Isolation of CD11b^+^ cells from adipose tissue

For isolation of CD11b^+^ adipose tissue macrophages (ATMs), adipose tissues from 8- to 12-week old male wild-type mice were collected and minced with scissors, before they were digested in a solution containing 1,5U/mL Collagenase D (Miltenyi Biotec, Germany), 2,4U/mL Dispase (Thermo Fisher Scientific) and 1mM CaCl_2_ for 45 minutes. Adipose tissues of 3-6 mice were pooled per sample and 6mL of digestion solution were used per sample. The resulting solutions were filtered through 100 μM and 40 μM cell strainer and were subjected to centrifugation at 600xg for 5 minutes. Afterwards, the supernatant was discarded and the pellet was used to perform magnetic separation of CD11b^+^ cells following the manufacturer’s instructions (Miltenyi Biotec GmbH, Germany). In brief, the cell pellet was re-suspended in 900 μl MACS buffer and incubated with 100 μl CD11b MicroBeads (Miltenyi Biotec GmbH, Germany) at 4°C for 15 min. After centrifugation, the cell pellet was re-suspended and the cell suspension, including the magnetically labeled CD11b^+^ macrophages, was applied to LS columns (Miltenyi Biotec GmbH, Germany). After washing, CD11b^+^ ATMs were collected from the LS column.

### Lipidomics of adipose tissue macrophages

Adipose tissue CD11b^+^ cells were enriched using magnetic isolated cell sorting (MACS® by Miltenyi Biotec GmbH, Germany). Lipidomic analysis was performed as described previously (*82*, *83*) utilizing the Lipidizer^TM^ platform from SCIEX. Briefly, lipids were extracted from cell pellets using an adjusted MTBE/methanol extraction protocol (*84*). Lipid extracts were concentrated and reconstituted in running buffer (10 mM ammonium acetate, dichloromethane (50): methanol (50)) and analyzed by a flow injection analysis-tandem mass spectrometry (FIA-MS/MS)-based method using an ultra-high pressure liquid chromatography system (Nexera X2, Shimadzu, Kyoto, Japan) coupled with a QTRAP® system (QTRAP® 5500; SCIEX) run in multiple-reaction-monitoring mode and equipped with a differential mobility spectrometer (DMS) interface operating with SelexION technology. Aquired raw data was processed and lipids were quantified using the Shotgun Lipidomics Assistant, a Python-based application (*82*). The samples were normalized on DNA content using dsDNA Kit, Broad Range (Thermo Fisher).

### Analysis of scRNA-seq data

Publicly available single-cell RNA-seq data of murine brown adipose tissue stromal vascular fractions (GSE160585) were analyzed using Seurat (*85*) (v4.0) in R (v4.3.1). Low-quality cells were excluded by retaining cells with >400 detected genes and >800 total transcripts. Data were normalized with LogNormalize and scaled using ScaleData. Highly variable genes were identified with the vst method (FindVariableFeatures, 2000 genes). Principal component analysis (PCA) was performed on the variable gene set, and the top 50 PCs were selected based on ElbowPlot. Clustering was carried out with FindNeighbors and FindClusters (resolution 0.2). Clusters were annotated by integration with published metadata and supervised curation based on marker gene expression. Functional module scores were computed using AddModuleScore with genes for lipid score: Lpl”, “Apoe”, “Lipa”, “Lipg”, “Lipe”, “Pnpla2”, “Lipe”, “Ldlr”, “Fabp4”, “Slc27a1” and Efferocytic score: *Axl, Mertk, Stab2, Trem2, Cd36, Scarf1, Timd4, Itgb3, Itgb5, Adgrb1, Gas6, Pros1, Thbs2, Mfge8, Nr1h3, Pparg, Nr1h2, Abca1, Pltp, Abcg5, Slc2a4, C1qb, C4b.* Wilcoxon rank-sum tests were applied to module scores between conditions, with false discovery rate (FDR) correction for multiple comparisons.

### *Ex vivo* Efferocytosis assays (ATMs)

After isolation of CD11b+ cells from adipose tissue, cells were counted and plated in a 48-well plate format at the highest possible density to ensure equal cell numbers across all samples (approximately between 50.000 cells/well to 200.000 cells/well). Cells were seeded in RPMI 1640 GlutaMAX^TM^ (Thermo Fisher Scientific) media containing 10% fetal bovine serum (FBS, Thermo Fisher Scientific), 10-20% L929-supplement (L929 cell supernatant in RPMI 1640 GlutaMAX^TM^ (*86*), and 1% Penicilin/Steptavidin (Thermo Fisher Scientific) and left to attach for approximately 2 h before CFSE (Thermo Fisher Scientific, C34554) labeled apoptotic thymocytes (aTs) were provided at 750.000 cells/well. ATMs were co-incubated with aTs for 1.5h at 37°C (Uptake) or on ice (Binding control) and then detached with cold PBS (Thermo Fisher Scientific no. C34554) after thorough washing. All cells were collected in round bottom 96-well plate and were stained for surface expression of LY6G, CD11b and F4/80 as well as with a viability dye (Invitrogen, L34961). For analysis of *ex vivo* efferocytosisis capacity of ATMs after NE injection, additionally, CX3CR1 and LY6C antibodies were added to the antibody mix. Finally, CD11b^+^F4/80^+^ ATMs were analyzed by flow cytometry for the presence of the aT-dye using FACSymphony™ A3 Cell Analyzer (BDBiosciences). The efferocytic index was defined as the difference between ATMs positive for the apoptotic cell dye (CFSE) at 37°C and 4°C. Data analysis was carried out using FlowJo software V10 (Tree Star).

### *In vitro* Efferocytosis assays (BMDMs)

BMDMs were differentiated from bone marrow precursors, from 6- to 14-week-old female mice. Briefly, femur and tibia were collected and cut open on one epiphysis and the hematopoietic pluripotent stem cells were isolated by centrifugation of the bones. The hematopoietic pluripotent stem cell homogenate was filtered over a 40 μm strainer and cells were seeded into RPMI 1640 GlutaMAX^TM^ (Thermo Fisher Scientific) containing 10% fetal bovine serum (FBS, Thermo Fisher Scientific), 10-20% L929-supplement (L929 cell supernatant in RPMI 1640 GlutaMAX^TM^ [323], and 1% Penicilin/Steptavidin (Thermo Fisher Scientific). During the propagation and differentiation of BMDMs, half of the media was changed at day 3. At day 5, cells were split and at day 7, the differentiated BMDMs were collected and seeded for the experiment. 0.33 × 10^6^ cells/wells were seeded into 24-well plates in the evening before the start of treatment. Treatments were carried out in the same RPMI media containing FBS, 10-20%, L929-supplement, and Penicilin/Steptavidin. Compounds used for treatment of BMDMs were diluted in either media or DMSO according to their properties and appropriate vehicle treatments were added to the controls. Norepinephrine was always prepared fresh and dissolved either in 0.9 NaCl (*in vivo*) or cell culture medium (*in vitro*). To assess the efferocytosis capacity of BMDMs, CFSE (Thermo Fisher Scientific no. C34554) labeled aTs were provided at a 5:1 ratio to BMDMs. BMDMs were co-incubated with aTs for 45 minutes at 37°C (Uptake) or on ice (Binding control) and then detached with cold PBS (Thermo Fisher Scientific no. C34554) after thorough washing. All cells were collected in round-bottom 96-well plates and were stained for surface expression of CD45, CD11b and F4/80 as well as with a cell viability dye (Invitrogen, L34961). Finally, CD11b^+^F4/80^+^ BMDMs were analyzed by flow cytometry for the presence of the aT-dye using a FACSymphony™ A3 Cell Analyzer (BDBiosciences). The efferocytic index was defined as the difference between BMDMs positive for the apoptotic cell dye (CFSE) at 37°C and 4°C. Data analysis was carried out using FlowJo software V10 (Tree Star).

### Preparation and staining of apoptotic thymocytes

For the isolation of thymocytes, one or more thymuses were harvested and smashed over a 40 µm strainer to obtain a single-cell solution. Cells were then pelleted and re-suspended in RPMI 1640 GlutaMAX^TM^ (Thermo Fisher Scientific) containing 5% fetal bovine serum (FBS, Thermo Fisher Scientific). Induction of apoptosis in thymocytes was done by incubating them overnight in 6mL of this low serum media. Apoptosis levels were verified using the FITC Annexin V Apoptosis Detection Kit (BioLegend no. 640914), following the manufacturer’s instructions. Next, aTs were stained with CFSE dye (Thermo Fisher Scientific no. C34554) for 20 minutes at 37°C at a concentration of 0.2 μL/mL and 10^8^ cells.

### Generation of NE-conditioned brown adipocyte supernatant

To generate NE-conditioned brown adipocyte supernatant, primary brown adipocytes were differentiated from stromal-vascular fractions (SVF) from BAT and WAT of wild-type mice (C57BL/6J, male, age 4-6 weeks) as described previously (*86*, *87*). In brief, adipose tissues were digested for 45 minutes using a solution containing 1.5% BSA (Sigma-Aldrich) and 600U/mL collagenase II (Sigma-Aldrich) in adipocyte isolation buffer (123 mM NaCl, 100 mM HEPES, 5 mM Glucose, 5 mM KCl, 1.3 mM CaCl2, pH 7.4). The obtained stromal vascular cells were subsequently cultured in DMEM GlutaMAX^TM^ media (4.5g/L Glucose, Thermo Fisher Scientific) supplemented with 10% newborn calf serum (Sigma-Aldrich, N4637-500ML), 1% penicillin and streptomycin (Thermo Fisher Scientific), 2.4 nM of insulin (Sigma-Aldrich) and 1µM of rosiglitazone (Cayman Chemicals). After 7 days of differentiation, mature adipocytes were treated with 1μM NE (Cayman Chemicals) and the resulting media was transferred in whole to BMDMs.

### Bulk RNA-sequencing

RNA isolations for bulk RNA sequencing were carried out as described for quantitative PCR and 200ng of RNA per sample were sent to BGI (BGI TECH SOLUTIONS, Hongkong) where RNA sequencing was performed.

For Bulk RNA sequencing of mature adipocytes (mAdipocytes) and SVFs, murine brown adipose of 8- to 12-week old male wild-type mice were were collected and minced with scissors, before they were digested in a solution containing 1,5U/mL Collagenase D (Miltenyi Biotec, Germany), 2,4U/mL Dispase (Thermo Fisher Scientific) and 1mM CaCl_2_ for 45 minutes. The resulting solutions were filtered through 100 μM and 40 μM cell strainer and were subjected to centrifugation at 600xg for 5 minutes. The upper fatty layer (mAdipocytes) as well as the pellet (SVF) were collected in 1 ml of TRIzol reagent (Thermo Fisher Scientific). Raw read count tables and sample metadata were imported into R (v4.3.1). Genes with zero counts across all samples were removed. Analyses were performed for two fractions separately: stromal vascular fraction (SVF) and mature adipocyte fraction (mAdipo). For each fraction (SVF and mAdipo), size factors were estimated and differential expression across temperature conditions were computed with DESeq2 (*88*).

For Bulk RNA sequencing of NE treated BMDMs, cells were differentiated and seeded as described earlier and and treated with 1µM NE or vehicle for 24h. Library preparation and transcriptome sequencing were performed using 100 bases/paired-end reads on BGI’s DNBSEQ Technology Platform. The mRNA sequencing with subsequent filtering (removal of adapters, low-quality reads, N reads and polyX) and quality control was performed on DNBSEQ Technology by BGI. For differential expression between controls and NE-treated BMDMs, reads for each sample were aligned to the mouse genome using STAR aligner (version 2.7.10b) and the resulting data was analyzed using the Dr. Tom web-based data visualization and analysis platform (Beijing Genomics Institute, BGI, accessed January 2025 (https://www.bgi.com/global/service/dr-tom).

For generation of PCA plots gene-level counts were generated from aligned RNA-seq reads (BAM files) using the summarizeOverlaps function from GENCODE vM36 and raw read count tables and sample metadata were imported into R (v4.3.1). Count data were normalized and variance-stabilized in DESeq2 and principal component analysis (PCA) was performed on VST-transformed counts. To assess group separation, PERMANOVA (adonis2, vegan R package) was conducted on Bray-Curtis distances calculated from transformed counts, using 20 permutations, with condition as the grouping variable.

### Quantitative PCR

Tissues or cell pellets were homogenized in 1ml of TRIzol^TM^ reagent (Thermo Fisher Scientific) using a Tissue Lyser type 3 (QIAGEN; 20 Hz for 2 × 3 min). 250 μL chloroform was added, and samples were mixed and centrifuged at 13,000 xg at 4°C for 15 min. The resulting supernatant was mixed with 600 μL of 70% ethanol. Further purification was performed by using NucleoSpin RNAII Kit (Machery&Nagel) according to the manufacturer’s instructions. Double-stranded DNA was digested using rDNase (Machery&Nagel). RNA content was measured using NanoPhotometer® N60 (IMPLEN, Germany) and subsequently 400 ng of RNA was transcribed to cDNA using High-Capacity cDNA Reverse Transcription Kit (Applied Biosystems). The reverse transcription PCR program was as follows: 1. 10 min, 25°C; 2. 120 min, 37°C; 3.5 s, 85°C. Gene expression was assessed in cDNA samples using Taqman® assays supplied as assays-on-demand^TM^ (Applied Biosystems) together with Taqman^TM^ Universal PCR MasterMix (Applied Biosystems) or by using custom primers from Eurofins together with SYBR™ Green Universal Master (Applied Biosystems). PCR reactions were performed by QuantstudioTM 5 Real-Time PCR Systems (Applied Biosystems) and cycling parameters were as followed: 1 cycle of 95°C for 10 min, 40 cycles of 95°C for 15 s then 60°C for 60 s. Gene copy number was calculated with the formula (10((Ct-35)/−3.3219)) and gene expression was always normalized for copy number of TATA-box binding protein (*Tbp*).

### Organelle proteomics

Organelle separation from BAT was carried out using male C57BL/6J mice, which were housed in virus-antibody-free facilities with unrestricted access to autoclaved food and water. For the experiments, BATs from 5 mice per group aged 12 weeks, housed under different conditions including room temperature, thermoneutrality (one week at 30°C), short cold exposure (4 hours at 4°C), and long cold exposure (1 week at 4°C), were used. Tissues were gently homogenized using a glass homogenizer on ice, and the lysates were loaded onto sucrose gradients. These gradients were prepared as per Klingelhuber et al. (2024) (*89*), using 6–7 mL of 20% sucrose solution followed by the addition of 50% sucrose solution, and stored at 4°C after mixing with the BioComp Gradient Master 108. For ultracentrifugation, pre-cooled equipment was used, and the gradients were centrifuged at 100,000 g for 3 hours at 4°C using a Beckman Optima L-70 ultracentrifuge with an SW41 rotor. Post-centrifugation, 24 fractions were collected from each tube in 0.5 mL increments and fractions 12 to 24 were diluted 1:1 with MilliQ water. Samples were mixed with 4 times the sample volume of ethanol precipitation buffer and stored at −20°C overnight.

Fractions 12–24 were diluted 1:1 with MilliQ water, followed by the addition of an ethanol precipitation buffer and stored at −20°C overnight. Proteomics involved sample preparation for LC-MS/MS analysis as described by Kulak et al. (2014) (*90*), with slight modifications. After centrifugation at 15,000 g for 15 minutes, the supernatant was discarded, and the remaining solvent was evaporated. Pellets were resuspended and lysed in SDC lysis buffer (2% sodium deoxycholate, 100 mM Tris-HCl, pH 8.5) at 95°C for 10 minutes, followed by sonication using a Diagenode Bioruptor® Plus. The protein concentration was determined by the bicinchoninic acid (BCA) assay (Thermo 23225), and 25 µg of protein were taken for further processing. Samples were treated with 10 mM tris(2-carboxyethyl)phosphine (TCEP) and 40 mM chloracetamide (CAA), incubated at 45°C for 10 minutes, and digested overnight with trypsin (Sigma t6567) and LysC (Wako 129-02541) at a 1:50 protein-to-enzyme ratio. After digestion, samples were acidified with 2% TFA in MS-grade isopropanol. Custom-made StageTips with three layers of SDB-RPS membranes were used for desalting and purifying the acidified samples. The peptides were eluted with an 80% ACN and 1.25% NH_4_OH buffer, dried in a vacuum centrifuge, and reconstituted in a loading buffer for LC-MS/MS analysis on an EASY-nLC 1200 system (Thermo Fisher Scientific) connected to an Orbitrap Exploris 480 Mass Spectrometer (Thermo Fisher Scientific) with injection via electrospray ionization (ESI), combined with a FAIMS Pro (Thermo Fisher Scientific). The peptides were separated on a 50 cm in-house packed HPLC column (75 µm innder diameter, packed with ReproSil-Pur C18-AQ 1.9µm resin, Dr. Maisch) using a binary buffer system (buffer A: 0.1% formic acid, buffer B: 80% acetonitrile, 0.1% formic acid), and a DIA mode was employed for data acquisition.

LC gradients started with 5% buffer B and increased to 45% over 45 minutes, followed by a washout at 95 % buffer B for 5 minutes. FAIMS compensation voltage was set to −50 V. MS1 scans (300 to 1,650 m/z, 45 ms maximum fill time, 300% normalized AGC target, R = 120,000 at 200 m/z) were followed by 66 unequally sized DIA windows, starting at a first mass of 120 m/z (22 ms fill time, 1,000% normalized AGC target, 30% normalized HCD collision energy, R = 15,00 at 200 m/z) in positive ion mode.

The raw data were analyzed using Spectronaut version 15.7 in directDIA mode against the human Uniprot FASTA database (2020), incorporating MaxLFQ for protein quantification. The analysis included data from the MaxQuant contaminant fasta file. Fractionation data were imported into C-COMPASS (Haas et al., bioRxiv, 2024), where samples were categorized by condition, replicates, and fractions. Default parameters and setting were applied for analysis. The datasets were filtered to include proteins identified in at least two replicates per condition, and missing values were replaced by zeros. Class Contribution (CC) values were calculated by averaging the outputs from neural network analysis of the three replicates, applying a 95% threshold to minimize false positives, and normalizing the values across compartments to ensure they summed to 1 per protein.

### Cytokine secretion assay on tissue explant

To measure cytokine secretion from iBAT and ingWAT, freshly isolated whole adipose tissues were collected and placed in DMEM GlutaMAX^TM^ media (4.5g/L Glucose, Thermo Fisher Scientific) supplemented with 10% fetal bovine serum (FBS, Thermo Fisher Scientific), 1% penicillin and streptomycin (Thermo Fisher Scientific). Afterwards medium was discarded and the explants were transferred to fresh medium and incubated for another 4 h at 37°C. Cytokines secreted into the media by iBAT explants were assessed using the LEGENDplex– Th1/Th2 Panel (8-plex) Assay (BioLegend no. 741054) following the manufacturers protocol. Data was collected using the FACS LSR and was analyzed via LEGENDplex software v8.0 (BioLegend).

### Extracellular flux analysis (Seahorse)

Oxygen consumption rate was assessed using a Seahorse XFe96 analyzer (Agilent Technologies). CD11b^+^ cells were resuspended in RPMI 1640 Glutamax^TM^ (Thermo Fisher Scientific) media supplemented with 10% fetal bovine serum (FBS, Thermo Fisher Scientific), 10% L929-conditioned medium, and 1% penicillin and streptomycin (Thermo Fisher Scientific). A total of 250,000 cells per well were seeded and allowed to adhere for approximately 3h. For the Mito stress test, media was replaced with Seahorse XF DMEM medium (pH 7.4) containing 10 mM glucose, 2 mM glutamin and 1 mM pyruvate. The assay was performed using 2 µM oligomycin, 2 µM FCCP (carbonylcyanide-p-trifluoromethoxyphenylhydrazone) and 1 µM rotenone/antimycin-A (all Sigma-Aldrich). For data normalization, cells were fixed with 4% PFA (paraformaldehyde) solution and stained with DAPI (Sigma-Aldrich). Fluorescence intensity was measured using a Tecan multimode plate reader (excitation: 350 nm; emission: 465 nm).

### Microscopy

Immunofluorescence imaging of efferocytosis assay. Efferocytosis assay was carried out using BMDMs as well as CFSE stained apoptotic thymocytes as described previosuly. After the assay BMDMs were stained with CD11b(APC-Cy7) and F4/80(AF700) and fixed with 2% paraformaldehyde for 20 min at room temperature. Fixed samples were placed on a 10 mm diameter cover glass, previously cleaned with a PlasmaCleaner (PDC-32G-2, Harrick Plasma) for 1 min.

All images were acquired using an inverted microscope (Leica Dmi8) equipped with an APO 10x/0.45 PH1, FL L 20x/0.40 CORR PH1, and APO 40x/0.95 objectives. Images were recorded with an ORCA-Flash4.0 Digital camera (Hamamatsu Photonics) using the MetaMorph Version 7.10.3.279 software (Molecular Device) and reconstituted by using Fiji (ImageJ) (*91*).

### Western blotting

Protein lysates were prepared by homogenizing adipose tissues using metal beads and a tissue lyzer (Qiagen). During this procedure, adipose tissues were kept in cold RIPA buffer ((10x (v/w), 20 mM Tris-HCl pH 7.4, 5 mM EDTA, 50 mM sodium chloride, 10 mM sodium pyrophosphate, 50 mM sodium fluoride, 1 mM sodium orthovanadate, 1 % NP-40, 0.1% SDS supplemented with Complete Mini Protease Inhibitor Cocktail Tablets (Roche) and phosphatase inhibitors (Sigma-Aldrich)). Lysates were centrifuged at 10.000 xg, for 5 min at 4°C and the resulting aqueous phase without the fatty layer was taken for protein determination using the Pierce™ BCA Protein Assay (Thermo Fisher Scientific). Protein samples were diluted in RIPA under addition of 4xNuPAGE sample buffer (Invitrogen) and 10x reducing agent (Invitrogen), so that 50 μg protein could be loaded per lane. Resulting samples were denatured for 10 min at 65°C. Samples were separated by SDS-PAGE on a 10% SDS-polyacrylamide Tris-glycine gels and were transferred to a nitrocellulose membrane (Amersham) overnight using a wet blotting system. Ponceau S (Sigma-Aldrich) staining was performed to ensure equal loading and membranes were cut according to the size of the proteins of interest. Membrane fragments that were to be stained for p563HSL were blocked in Roti®Block (Carl Roth, 1:10 diluted with aqua dest) for 1h at RT, while all other membrane fragments were blocked in 5% milk (Millipore) in TBST (0.2 M Tris, 1.37 M sodium chloride, 0.1 % (v/v) Tween 20). After washing with TBST, membrane fragments were incubated with primary antibody in 5% Bovine Serum Albumin (BSA, Sigma-Aldrich) +0.02% NaAzid in TBST overnight at 4°C. Primary antibodies used were (BAX (Cell signaling, #2772, 1:1000); rabbit-anti-gTubulin (Abcam, TUBG1, ab179503, 1:2000); mouse-anti-UCP1 (R&D Systems, MAB6158, 1:500); rabbit-anti-p563HSL (Cell signaling, 413S, 1:1000); rabbit-anti-HSL (Cell signaling, 4107S, 1:1000); rabbit-Phospho-PKA Substrate (Cell signaling, 9621, 1:1000); rabbit-anti-ADRB3 (Abcam, ab94506, 1:1000) rabbit-anti-Tyrosine Hydroxylase (TH, Abcam, ab137869, 1:1000); rat-anti-MAC2 (Santa Cruz, sc23938, 1:1000). After another round of thorough washing with TBST, membrane fragments were incubated with secondary antibody in 5% milk in TBST for 1h at RT. Secondary antibodies used were HRP-conjugated goat anti-rabbit (Cell Signaling, #7074, 1:5000); HRP-conjugated rabbit anti-mouse (CiteAb Ltd, P0161, 1:5000) or HRP-conjugated donkey anti-rat (Jackson ImmunoResearch, AM_23040641, 1:5000). After more rounds of washing in TBST, membrane fragments were developed on Amersham Imager600 (GE Healthcare) using either luminol (Sigma-Aldrich) or SuperSignal™ West Femto Maximum Sensitivity Substrat (Thermo Scientific™ 34096X4). Quantification of the bands was done in Image Studio Lite (version 5.2.5) and protein expression was normalized as indicated in the respective Figures. After quantification some membrane fragments were stripped using incubation with 1x Restore™ Fluorescent Western Blot Stripping Buffer (Thermo Fisher Scientific) and reincubated with another primary antibody to determine the levels of a second protein of interest. Quantification of results was done using ImageJ Version 1.54g. Protein expression was normalized as indicated in the respective Figures and the mean of the control group was set to 1.

### Indirect calorimetry

For studies on systemic energy expenditure, mice were put in metabolic cages (Sable Systems Europe GmbH, Germany) and the system was operated according to the manufactures guidelines. Mice were acclimated to the metabolic cages for at least 2 days at 28°C prior to recording. 28°C was used as the standard thermoneutral temperature and change of temperature occurred at 7 am and the light phase was from 7 am – 7 pm. During the experiment, consumption of O_2_, production of CO_2_, respiratory exchange ratio (RER), energy expenditure, food intake, water intake, body core temperature and activity were measured.

### Ex vivo Lipolysis assay

For ex vivo Lipolysis assays 2-3 small, freshly isolated iBAT pieces of around 20mg were weighed and placed in 200µl of DMEM, high glucose, HEPES, no phenol red (Thermo Fisher Scientific) supplemented with 0.2% fatty acid free BSA (Capricorn Scientific) for 1h at 37 °C. Afterwards, media was discarded and explants were transferred to fresh media containing FA-free BSA and incubated at 37 °C for 4 h. To stimulate lipolysis, explants were transferred to medium supplemented with an additional 1µM NE (Cayman Chemicals). NEFA release into the media was analyzed by using a colorimetric assay (NEFA-HR(2) Assay, FUJIFILM). Results were normalized to the weight of the respective explants and results from one mouse were averaged to equal one biological replicate.

### Blood and plasma parameters

After heart blood collection, whole blood was centrifuged at 4°C for 10 min at 10.000 xg for 5 min. The plasma was then transferred to a new reaction tube and was stored at −80°C if it was not directly analyzed. For plasma Triglyceride and cholesterol determination, 5μL of plasma was pipetted in duplicates into 96-wells plates and concentrations were determined using commercial kits following the manufacturer’s instructions (DiaSys Diagnostic Systems). For plasma non-esterified fatty acids (NEFA) concentrations 20 μL plasma was pipetted in duplicates into a 96-well plate and concentrations were determined using the commercial kit NEFA-HR(2) Assay by FUJIFILM.

### Histology – Immunohistochemistry

For histological analysis, tissues were fixed in 3.7% formaldehyde (diluted in PBS) and embedded in paraffin. 4μm paraffin sections were made, deparaffinized and subsequently rehydrated in xylene and descending ethanol series. For UCP1 staining slides were cooked in 10 mM citrate buffer (pH 6.0) for 30 minutes for antigen retrieval and left in the buffer for an additional 30 min to cool down. After washing, slides were blocked with 3% BSA in PBS for 1h at RT. For UCP1 IHC stainings we used rabbit-anti-UCP1 (Abcam, ab10983) in a dilution of 1:500 in 3% BSA (Sigma) and Horseradish peroxidase (HRP) coupled goat anti-rabbit (Cell Signaling, #7074) was used as secondary antibody. After addition of DAB-Substrate (Abcam) and Hematoxylin (Sigma-Aldrich) counterstaining slides were sealed using Eukitt (O.Kindler GmbH). For Hematoxylin and Eosin staining of BAT sections were stained witth Hematoxylin (Sigma-Aldrich) for 10 minutes and Eosin (Sigma-Aldrich) for 1 minute before rehydration and mounting. Images were taken using a NikonA1 Ti microscope equipped with a DS-Fi-U3 brightfield camera and quantified using Image J Version 1.54g.

### Statistics

Data are presented as mean ± SD (Standard deviation). Graphs as well as statistical analysis were made in Graph Pad Prism (Version 10.0). Outliers were identified for each graph using the ROUT method with a Q cut-off value of 1%. For all experiments with n<15 non-parametric statistical tests were performed and for experiments with n>15 normality was tested via D’Agostino and Pearson test before using the adequate statistical test. For analysis of lipidomics data unpaired ttests were used. For Figure 3, if repeated Measures two-way ANOVA followed by Šidák’s multiple comparison test was performed, comparisons were only made between control and the paired NE treated samples. Sample sizes, p values, as well as the statistical tests implemented, can be found in the figures or the figure legends. A p-value <0.05 was considered significant. No statistical method was used to predetermine sample sizes.

## Supporting information

Meyer et al. Suppl. Fig.

## Supplementary Materials

Supplementary figures S1 to S7

Supplementary Tables 1 and 2

## Acknowledgments

We thank Nicola Gagliani (University Medical Center Hamburg-Eppendorf, Hamburg, Germany) for providing helpful feedback and valuable discussions. We thank Alexander Bartelt and Imke Lemmer (Technical University of Munich, Germany) for performing and supporting the mouse experiments related to the proteomic analysis. We thank Anke Baranowksy and Johannes Keller (University Medical Center Hamburg-Eppendorf, Hamburg, Germany) for providing *Adrb2*^+/+^ and *Adrb2*^−/−^ mice and we thank Carla Rothlin (Yale University School of Medicine, New Haven, USA) for proviving *Csf1r*^Cre^*Axl*^f/f^*Mertk*^f/f^ mice. Schematic representation in fig. S4 was created with BioRender.com.

## Author contributions

Conceptualization: SM, LB, AW

Methodology: SM, JW, IL, TLL, LE, SL, JohH, DH, LA

Investigation: SM, CS, PJS, NK, JH, LA, LB, AW

Funding acquisition: LB, AW

Supervision: LB, AW

Writing – original draft: SM, LB, AW

Writing – review & editing: CS, PJS, NK, JH, LA, LB, AW

## Data and materials availability

All data needed to evaluate the manuscript are present in the main text or the supplementary materials. Sequencing fastq files and processed data will be made available upon revision. *Csf1r*^Cre^*Axl*^f/f^*Mertk*^f/f^ mice are available from Carla Rothlin (Yale University School of Medicine, New Haven, USA) under a material agreement with Yale University and a completed material transfer agreement.

