## Supplementary material for "Sympathetic signaling directs macrophage efferocytosis in thermogenic adipose tissue": Meyer et al. Suppl. Fig.

**fig. S1-S7**

**Table 1-2**

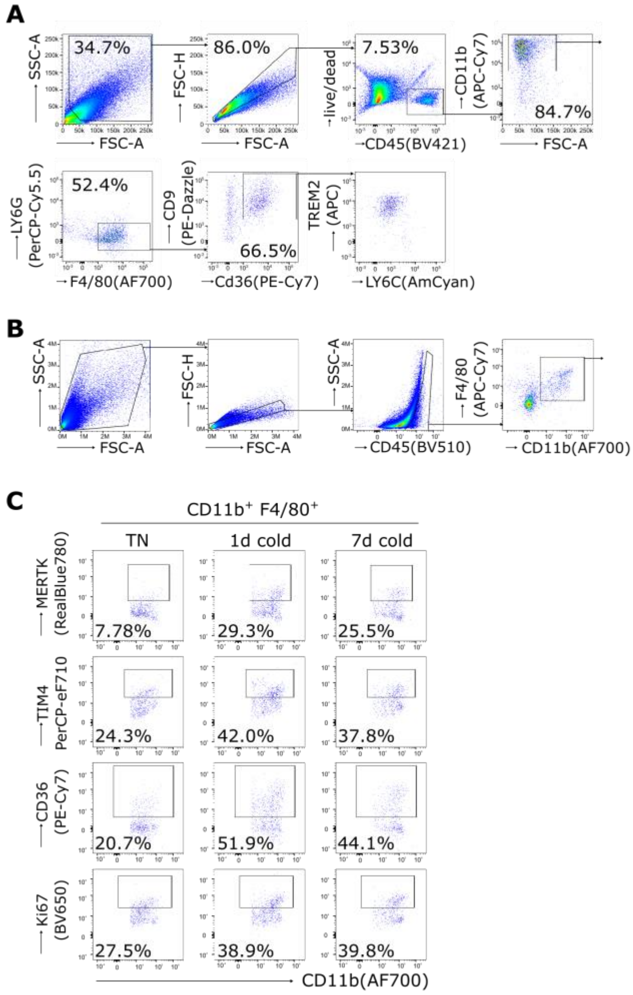


**fig. S1. iBAT macrophages during cold exposure.** (**A**) Gating strategy showing CD36$^+$CD9$^+$CD11b$^+$F4/80$^+$LY6G$^{low}$ macrophages in interscapular BAT (iBAT) of mice that were exposed to cold (6 °C) for 1 day; Gating strategy (**B**) as well as representative dot plots (**C**) for efferocytosis receptor expression in iBAT macrophages isolated from mice kept at thermoneutrality (TN, 30°C) or mice exposed to cold (6°C) for 1 day or 7 days.

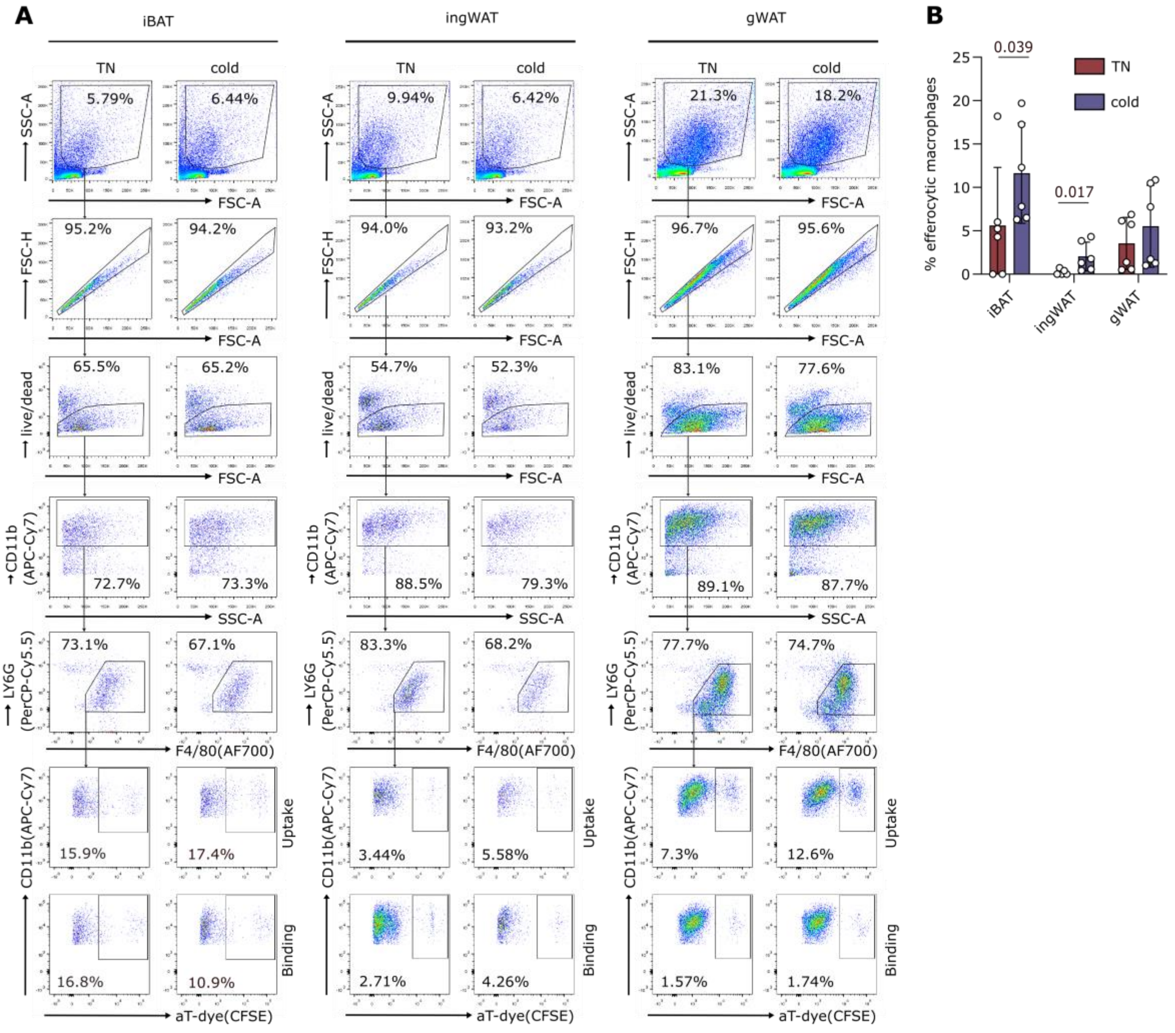


**fig. S2. Efferocytosis capacity of iBAT and WAT macrophages following cold exposure.** Gating strategy (**A**) and pooled results (**B**) for *ex vivo* efferocytosis capacity of Live$^{+}$CD11b$^{+}$LY6G$^{int}$F4/80$^{+}$ adipose tissue macrophages isolated from interscapular BAT (iBAT), inguinal WAT (ingWAT) or gonadal WAT (gWAT) of mice kept at thermoneutrality (TN, 30°C) exposed to cold (1d 6°C). Each dot represents a pool of 3 mice. n=6. Mean±SD. Mann-Whitney tests. One outlier was detected (ROUT method, Q=1%) and excluded from the corresponding analysis.

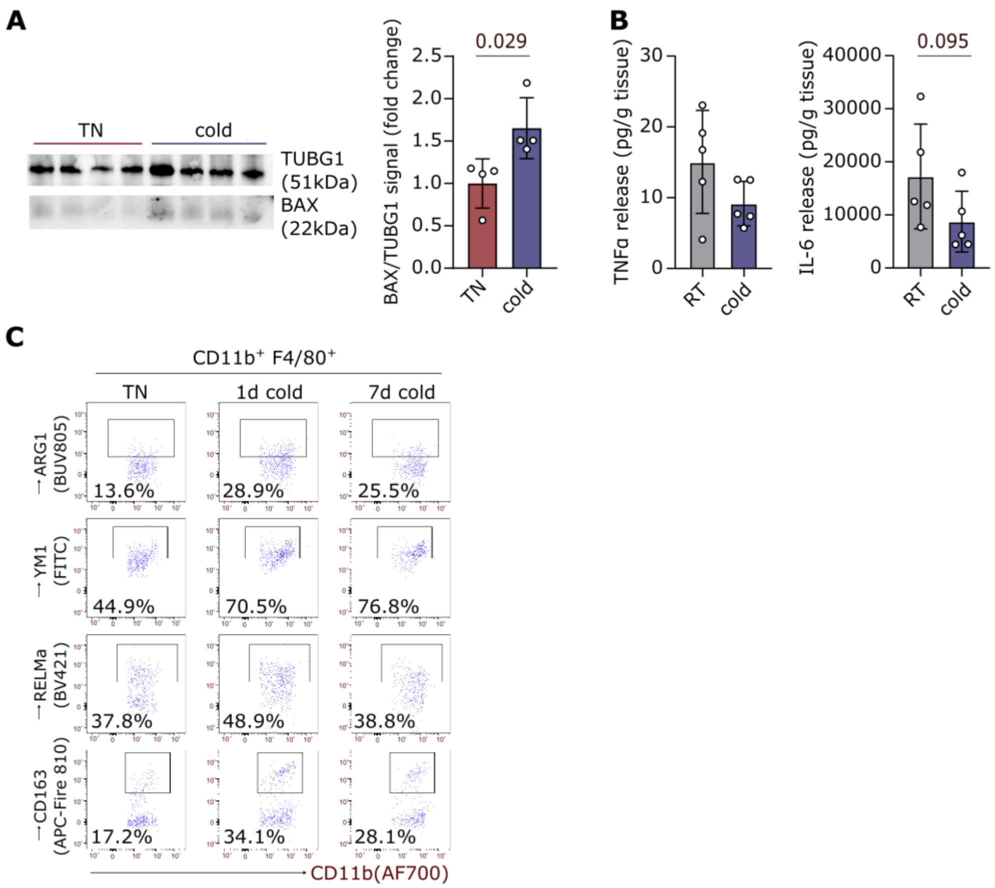


**fig. S3. Mice exposed to cold have higher BAX protein expression in iBAT along with a more anti-inflammatory immune milieu.** (**A**) BAX protein levels interscapular BAT (iBAT) of mice kept at thermoneutrality (TN, 30°C) or exposed to cold (6°C) for 1 day. Each dot represents one mouse. n=4. Mean±SD. Mann-Whitney test; (**B**) TNFα and IL-6 levels in supernatants of iBAT explants isolated from mice kept at thermoneutrality (TN) or room temperature (RT) assessed by LEGENDplex analysis after 4h of incubation. Each dot represents one mouse. n=5. Mean±SD. Mann-Whitney tests; Gating strategy (**C**) Representative dot plots for immune phenotyping of iBAT macrophages isolated from mice kept at thermoneutrality or mice exposed to cold for 1 day or 7 days.

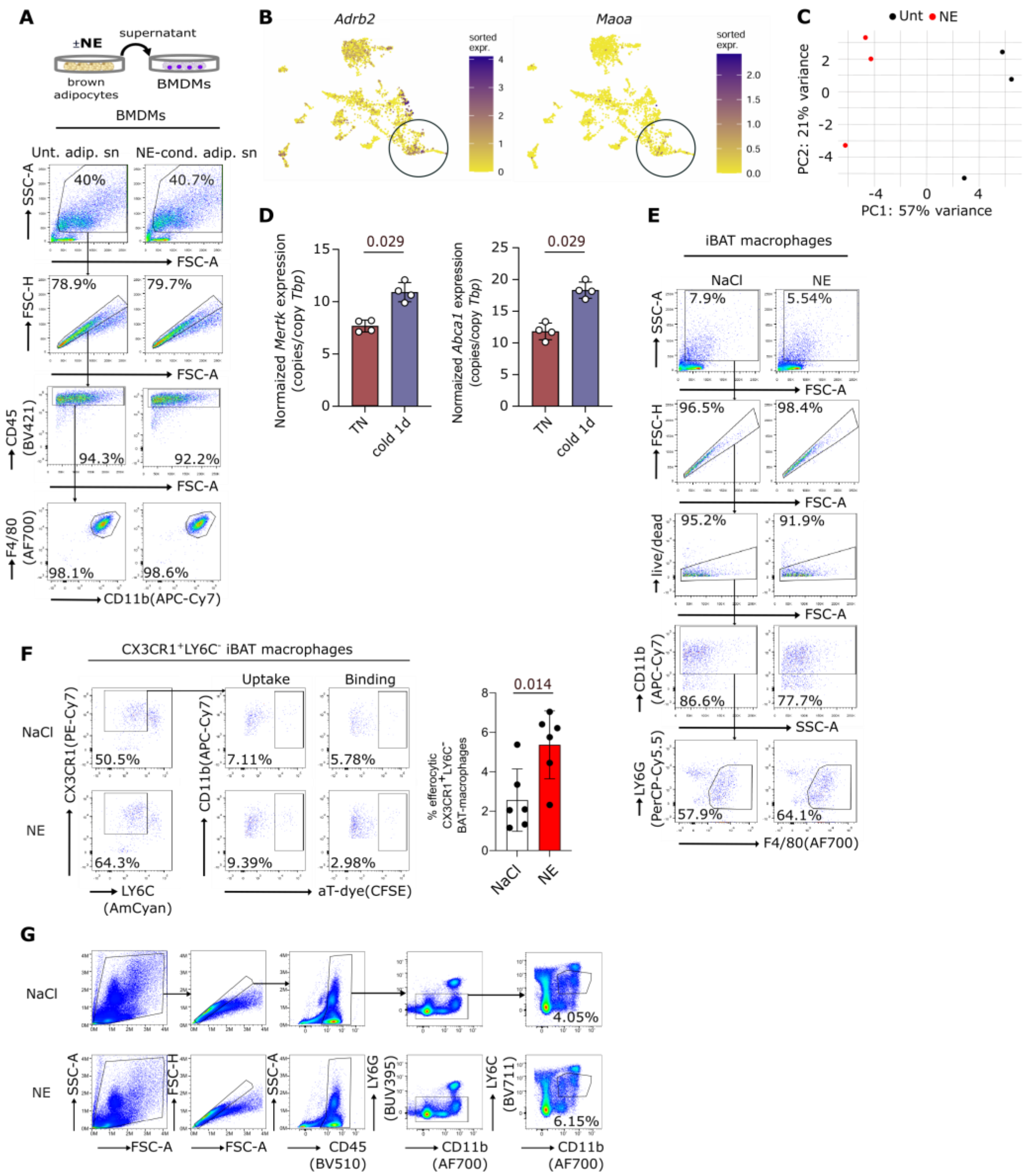


**fig. S4. NE sensing induces macrophages efferocytosis *in vitro* and *in vivo*.** (**A**) Gating strategy used to analyze of bone marrow-derived macrophages (BMDMs); (**B**) *Adrb2* and *Maoa* expression among all macrophages identified in single-cell sequencing data of murine BAT by Shamsi et al. 2021 *Nat. Metab*. (GSE160585); (**C**) PCA plot of untreated BMDMs, or BMDMs treated with 1 $\mu$M norepinephrine (NE) for 24h. Each dot represents one mouse. n=3. p=0.095, PERMANOVA; (**D**) Normalized *Mertk* and *Abca1* expression in CD11b$^{+}$ cells from brown adipose tissue of mice kept at thermoneutrality (TN) or exposed to cold (6°C) for 24h. Each dot represents a pool of 4-5 mice. n=4. Mean±SD. Mann-Whitney tests; (**E**) Gating strategy for analysis of murine intercapular BAT (iBAT) macrophages following subcutaneous injection with NaCl or norepinephrine (NE) at room temperature at 1mg/kg bodyweight twice, 12h and 15 minutes before analysis. Gating strategy (**E**) and pooled results (**F**) for *ex vivo* efferocytosis capacity of Live$^{+}$CD45$^{+}$CD11b$^{+}$F480$^{+}$CX3CR1$^{+}$LY6C$^{-}$ iBAT macrophages that were CD11b$^{+}$ isolated from mice injected subcutaneously with NaCl or norepinephrine (NE) at 1 mg/kg bodyweight twice, 12h and 15 minutes before analysis us. Injection was done in the interscapular region. Each dot represents a pool of 3 mice. n=6. Mean±SD. Mann-Whitney test. (**G**) Gating strategy to asses efferocytosis receptor expression in CD45$^{+}$LY6G$^{low}$CD11$^{+}$LY6C$^{int}$ non-classcical blood monocytes in mice after injection with NaCl or NE at 1mg/kg bodyweight twice, 12h and 15 minutes before analysis;

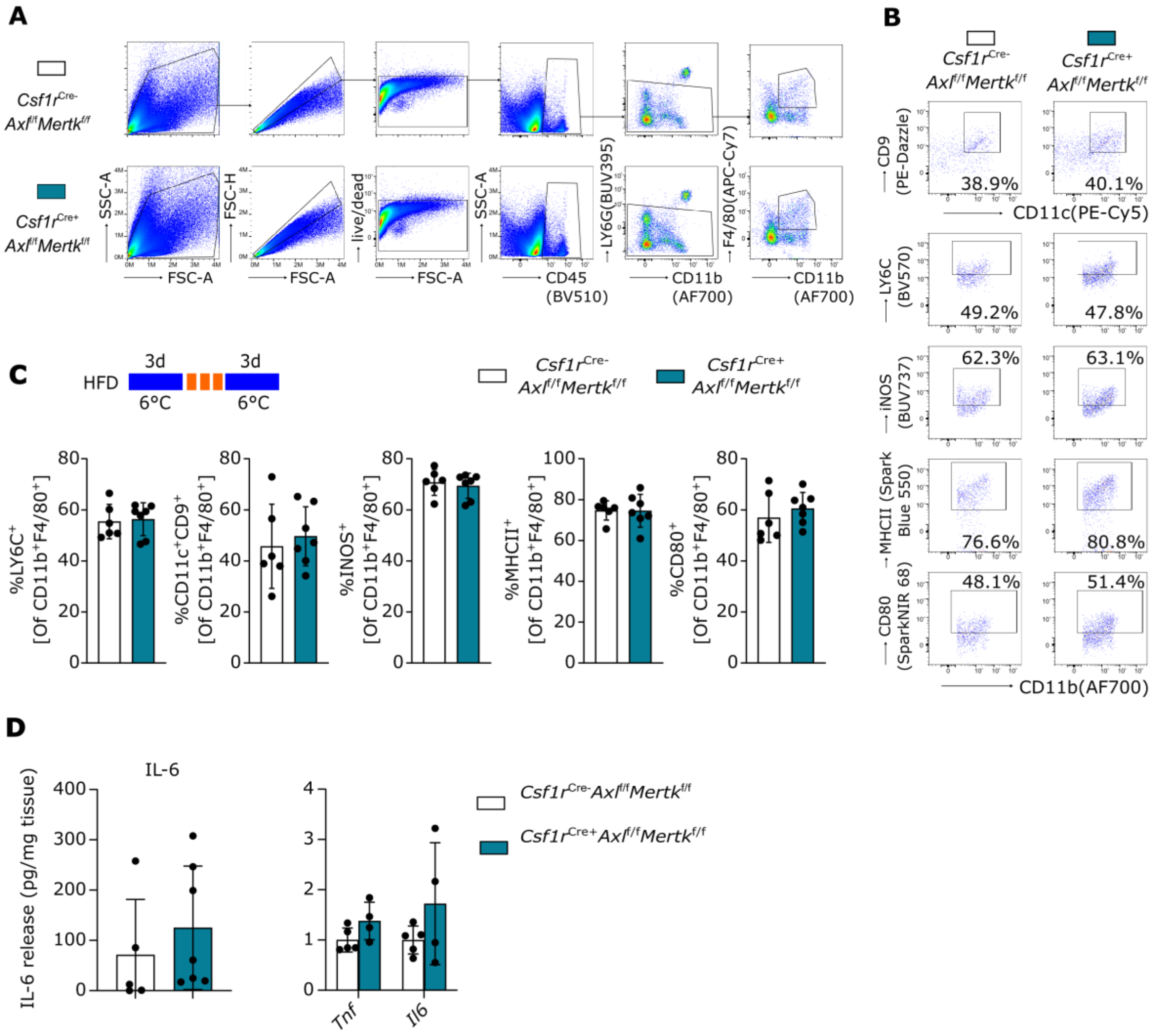

**fig. S5. Immune profiling of iBAT macrophages from mice fed with HFD and subsequent ly exposed to repeated cold cycles.** (**A**) Gating strategy, (**B**) representative dot plots and (**C**) results, comparing the inflammatory status of Live$^{+}$CD45$^{+}$LY6G$^{low}$CD11b$^{+}$F4/80 interscapular BAT (iBAT) macrophages between *Csf1r*$^{Cre-}$ *Axl*$^{f/f}$ *Mertk*$^{f/f}$ and *Csf1r*$^{Cre+}$ *Axl*$^{f/f}$ *Mertk*$^{f/f}$ mice after HFD and repeated cold exposure. n=6/7. Mean±SD. Each dot represents one mouse; (**D**) Secretion of IL-6 by iBAT explants of *Csf1r*$^{Cre-}$ *Axl*$^{f/f}$ *Mertk*$^{f/f}$ and *Csf1r*$^{Cre+}$ *Axl*$^{f/f}$ *Mertk*$^{f/f}$ mice into cell culture medium after 4h, as assessed by LEGENDplex analysis as well as expression of *Tnf* and *Il6* in iBAT, analyzed by qPCR. For both experiments, iBATs were collected after HFD feeding and subsequent repeated cold exposure as for ( A-C). Each dot represents one mouse. n=5/7 and n=4/5 respectively. Mean±SD;

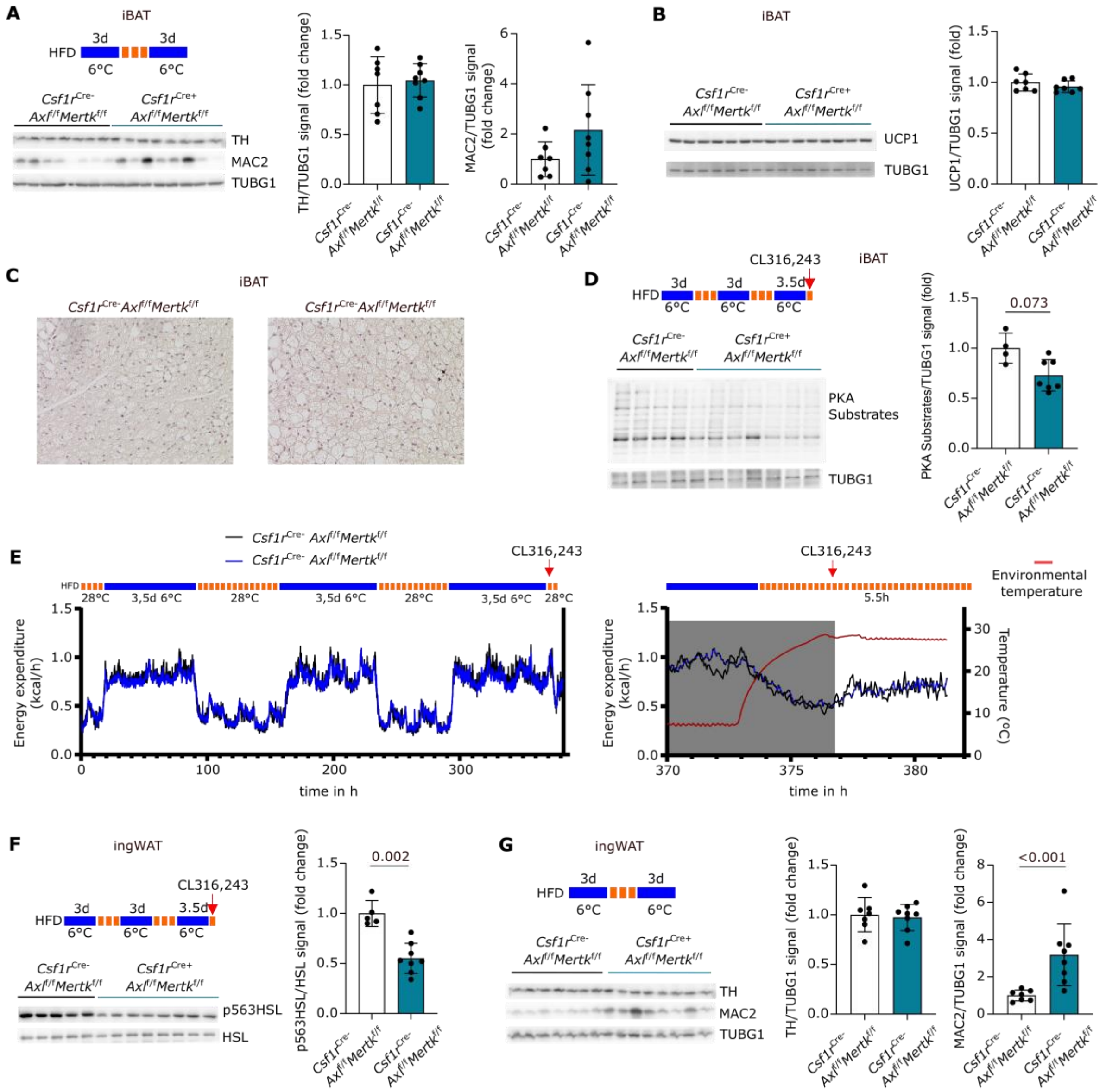

**fig. S6. *Axl* and *Mertk* deletion in macrophages has minor impact on iBAT thermogenic function.** (**A**) Tyrosine hydroxylase (TH) and MAC2 expression in interscapular BAT (iBAT) of *Csf1r*$^{Cre-}$ *Axl*$^{f/f}$ *Mertk*$^{f/f}$ and *Csf1r*$^{Cre+}$ *Axl*$^{f/f}$ *Mertk*$^{f/f}$ mice after HFD feeding and subsequent repeated cold exposure. n=7/8. Mann-Whitney tests; (**B**) UCP1 levels in iBAT of *Csf1r*$^{Cre-}$ *Axl*$^{f/f}$ *Mertk*$^{f/f}$ and *Csf1r*$^{Cre+}$ *Axl*$^{f/f}$ *Mertk*$^{f/f}$ mice after HFD feeding and subsequent repeated cold exposure as in A). n=7; **(C)** Representative sections from hematoxylin and eosin staining of iBAT from *Csf1r*$^{Cre-}$ *Axl*$^{f/f}$ *Mertk*$^{f/f}$ and *Csf1r*$^{Cre+}$ *Axl*$^{f/f}$ *Mertk*$^{f/f}$ mice treated as in (A). (**D**) PKA substrates levels in iBAT of *Csf1r*$^{Cre-}$ *Axl*$^{f/f}$ *Mertk*$^{f/f}$ and *Csf1r*$^{Cre+}$ *Axl*$^{f/f}$ *Mertk*$^{f/f}$ mice after HFD feeding and, repeated cold exposure and injection of CL316,243 five hours before organ harvest. n=4/7. Mann-Whitney tests. (**E**) Energy expenditure of *Csf1r*$^{Cre-}$ *Axl*$^{f/f}$ *Mertk*$^{f/f}$ and *Csf1r*$^{Cre+}$ *Axl*$^{f/f}$ *Mertk*$^{f/f}$ mice after HFD feeding during repeated cold exposure and after injection of CL316,243 at 1 mg/kg bodyweight. n=5/8; (**F**) p563HSL levels in iBAT of *Csf1r*$^{Cre-}$*Axl*$^{f/f}$ *Mertk*$^{f/f}$ and *Csf1r*$^{Cre+}$*Axl*$^{f/f}$ *Mertk*$^{f/f}$ mice treated as in (E). n=5/8. Mann-Whitney test; (**G**) TH and MAC2 in ingWAT of *Csf1r*$^{Cre-}$ *Axl*$^{f/f}$ *Mertk*$^{f/f}$ and *Csf1r*$^{Cre+}$ *Axl*$^{f/f}$ *Mertk*$^{f/f}$ mice after HFD feeding and subsequent repeated cold exposure. n=7/8. Mann-Whitney tests; In this Figure each point corresponds to one mouse and all data are displayed as Mean±SD. Protein levels were normalized to the mean expression in *Csf1r*$^{Cre-}$*Axl*$^{f/f}$ *Mertk*$^{f/f}$ mice.

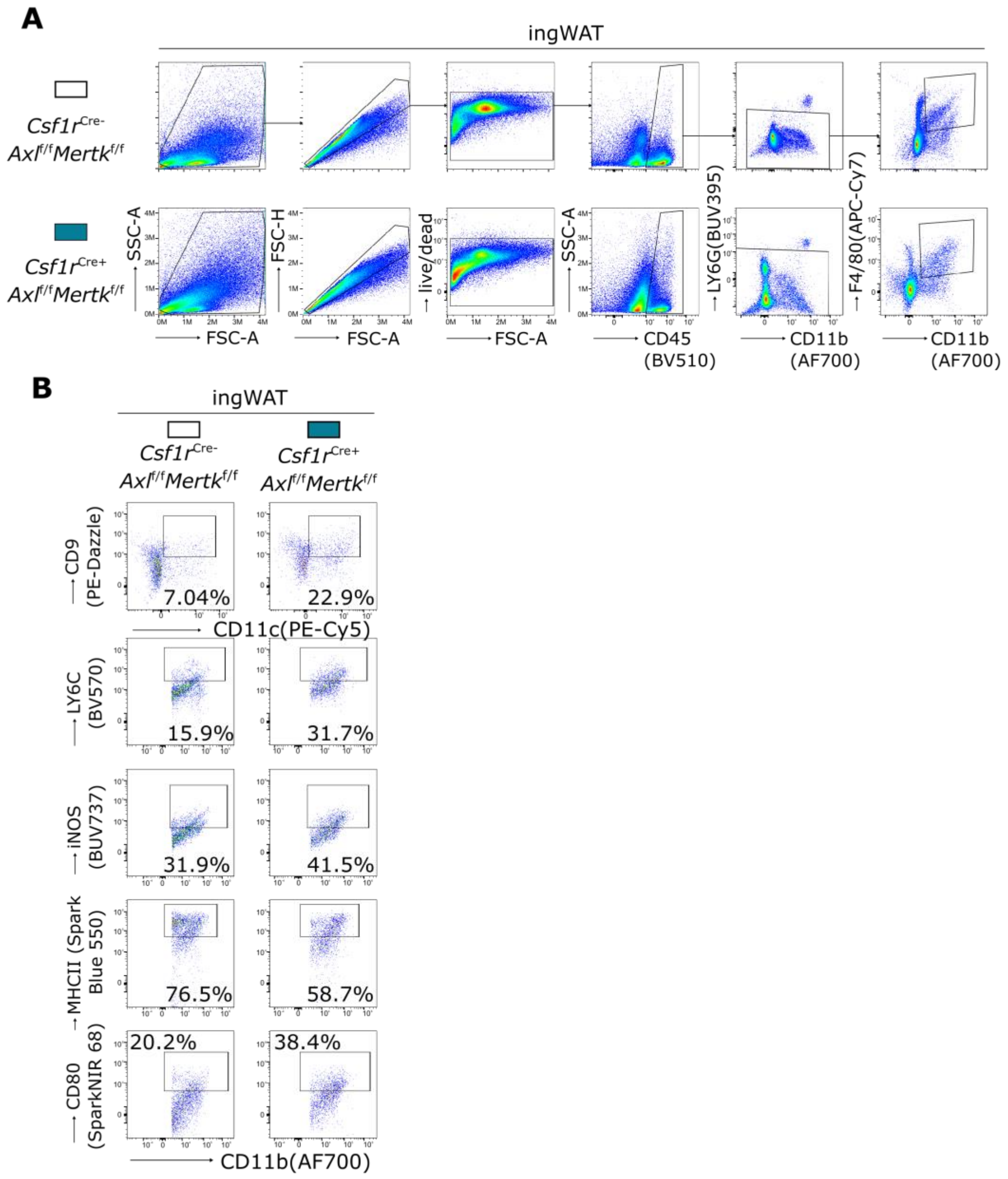


**fig. S7. ingWAT macrophages from *Csf1r*$^{cre+}$ *Axl*$^{f/f}$ *Mertk*$^{f/f}$ mice show a more inflammatory phenotype after HFD feeding and subsequent repeated cold exposure as compared to ingWAT macrophages from *Csf1r*$^{cre-}$ *Axl*$^{f/f}$ *Mertk*$^{f/f}$ mice.** (**A**) Gating strategy and (**B**) representative dotplots for flow cytometric analysis, assesing the inflammatory status of Live$^+$CD45$^+$LY6G$^{low}$CD11b$^+$F4/80$^+$ ingWAT macrophages isolated from *Csf1r*$^{cre-}$ *Axl*$^{f/f}$ *Mertk*$^{f/f}$ and *Csf1r*$^{cre+}$ *Axl*$^{f/f}$ *Mertk*$^{f/f}$ mice after HFD and subsequent repeated cold exposure as indicated in Fig. 5A.
